# Evaluation of chromosome structure modelling tools in bacteria

**DOI:** 10.1101/2023.10.26.564237

**Authors:** Tong Liu, Qin-Tian Qiu, Kang-Jian Hua, Bin-Guang Ma

**Affiliations:** Hubei Key Laboratory of Agricultural Bioinformatics, College of Informatics, Huazhong Agricultural University, Wuhan 430070, China

**Keywords:** Prokaryotes, Chromatin interaction, Hi-C, Chromosome modeling, Algorithm evaluation

## Abstract

The three-dimensional (3D) structure of bacterial chromosomes is crucial for understanding chromosome function. With the growing availability of high-throughput chromosome conformation capture (3C/Hi-C) data, the 3D structure reconstruction algorithms have become powerful tools to study bacterial chromosome structure and function. It is highly desired to have a recommendation on the chromosome structure reconstruction tools to facilitate the prokaryotic 3D genomics. In this work, we review existing chromosome 3D structure reconstruction algorithms and classify them based on their underlying computational models into two categories: constraint-based modeling and thermodynamics-based modeling. We briefly compare these algorithms utilizing 3C/Hi-C datasets and fluorescence microscopy data obtained from *Escherichia coli* and *Caulobacter crescentus*, as well as simulating datasets. We discuss current challenges in the 3D reconstruction algorithms for bacterial chromosomes, primarily focusing on software usability. Finally, we briefly prospect future research directions for bacterial chromosome structure reconstruction algorithms.

## Introduction

The three-dimensional (3D) conformation of chromosomes reveals important insights into how chromosome structure and function are related, such as gene regulation [1]. With the advancement of various biotechnologies, especially high-throughput sequencing, researchers have more tools to study the organizational and functional features of chromosomes. In 2002, the chromosome conformation capture (3C) technique opened a new avenue for studying the 3D conformation of chromosomes [2]. This technique has been further improved to enable the study of different types of DNA interactions: circular chromosome conformation capture (4C) [3] for one-to-many interactions; carbon-copy chromosome conformation capture (5C) [4] for many-to-many interactions; and Hi-C (high-throughput/resolution chromosome conformation capture) [5], which combined 3C with high-throughput sequencing technology, allows the study of all-to-all chromosome fragment interactions at the genome level. The later improved digestion-ligation-only (DLO) Hi-C technology greatly reduces background noise and experimental cost [6].

The sequencing data obtained from 3C/Hi-C experiments can be mapped to the genome to identify the positions of both ends of the DNA fragments. Then, the numbers of interactions between all fragments of the chromosome (comprising the interaction matrix) can be calculated. The number in the matrix represents the probability (intensity) of interaction between fragments. By analyzing the features of interaction matrix, it is found that the spatial conformation of chromosome is not random, and there is a phenomenon that certain fragments have significantly stronger interactions among themselves than with other fragments. These fragments are called TADs (topologically associating domains) [7]. TADs have been observed in different species [8-10].

One goal of 3D genomics is to construct the spatial structure of chromosomes by determining the spatial coordinates of each DNA fragment or even of each nucleotide. Currently, methods for chromosome modeling are broadly classified into two categories (**Figure 1**): thermodynamics-based modeling and constraint-based modeling [11]. Thermodynamics-based modeling methods employ polymer physics theory and/or numerical simulation to characterize the behavior of chromatin fiber. In this approach, chromatin fibers are conceptualized as a series of polymer units, and various factors, including inter-unit interaction (such as electric force) and conformational energy, are taken into account to determine the 3D chromosome structure [12]. This approach often uses Brownian dynamics and Monte Carlo simulation to address complex issues involving chromatin unit (e.g., nucleosome) diffusion and movement within the cell nucleus [13]. Due to the involvement of energy optimization, this method is effective for constructing models with a small number of fragments. If the number of fragments is too large, computational speed becomes a bottleneck. Constraint-based modeling methods emphasize the utilization of experimental data or known chromosome structural information to guide the inference of chromosome structure. An increasing amount of interaction data obtained through 3C/Hi-C experiments provide constraint information on the spatial structure of chromosome. The algorithms for constraint-based modeling come in many varieties and their implementation can be roughly divided into three stages (**Figure 2**): transforming the interaction frequency matrix (IF matrix) into an expected spatial distance matrix, defining the objective function and constraint parameters, and optimizing the objective function to obtain the final conformation [14]. Furthermore, these two categories of modeling methods are not entirely mutually exclusive, as some algorithms appropriately combine characteristics from different methods. For instance, Yildirim et al. incorporated distance restraints from Hi-C interaction frequency into their plectonemic supercoiling model for *Caulobacter crescentus*, and generated an ensemble of models with base pair resolution [15].

**Figure 1.**
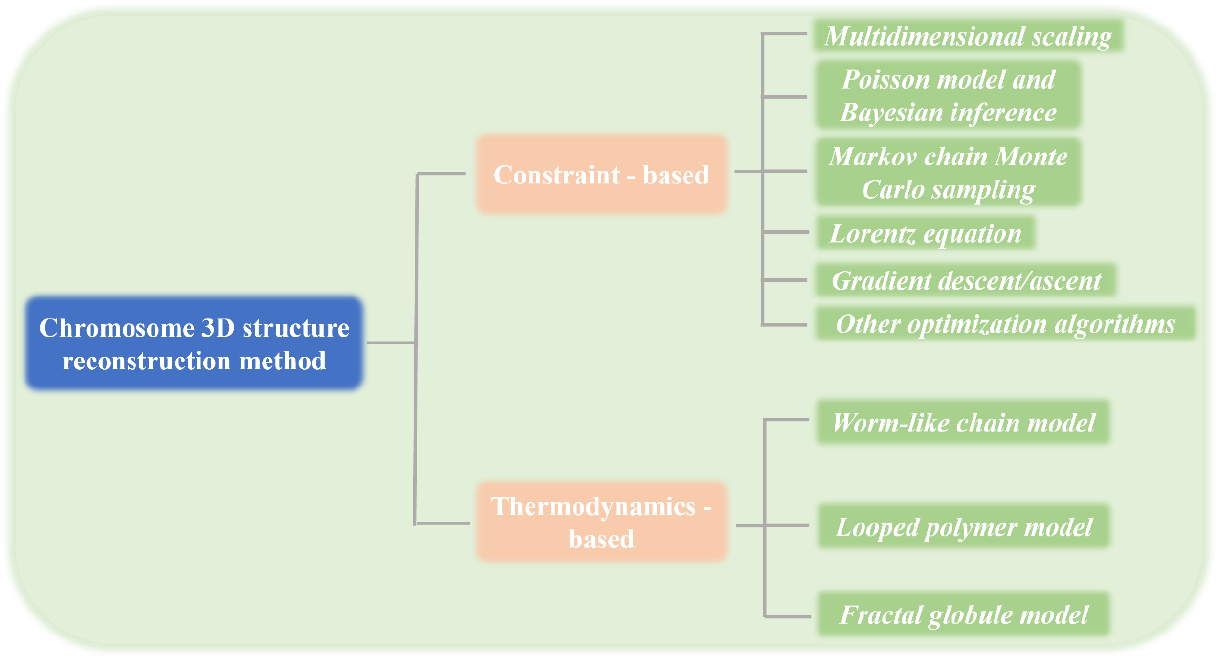
Classification of chromosome 3D structure reconstruction methods.

**Figure 2.**
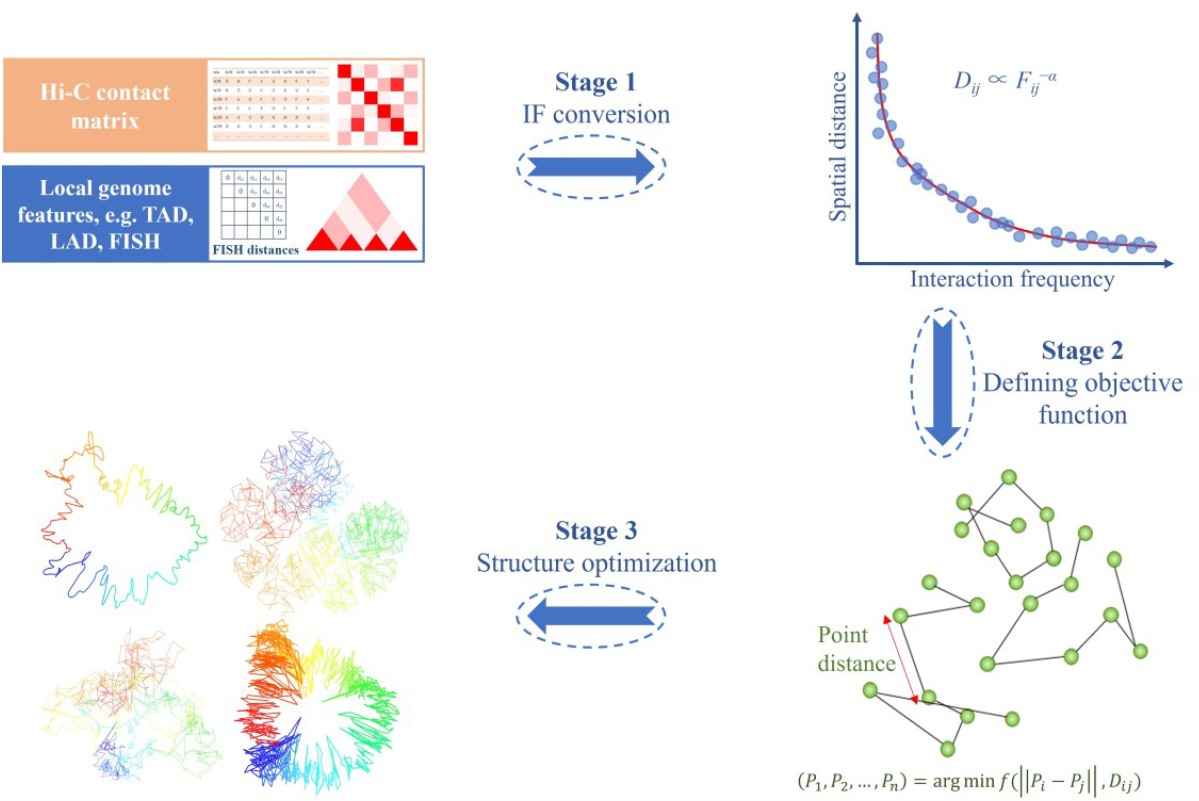
The general process of constraint-based modelling for chromosome 3D structure reconstruction. Stage 1: Convert the input data, usually Hi-C contact matrix or local genomic features, into distance matrix. Stage 2: Define the objective function and constraint parameters. Stage 3: Optimize the structure to obtain the final conformation.

The 3C/Hi-C technology was initially invented to investigate the 3D chromosome structure in eukaryotic organisms, yielded significant insights [5, 16, 17]. Over the past decade, it has also found extensive application in prokaryotic organisms, significantly advancing our understanding of the dynamic folding and interactions within bacterial genomes. In 2011, Umbarger et al. used 5C technology to produce an interaction map of *C. crescentus* with a resolution of 13kb, and they used the integrated modeling platform (IMP) to construct its model [18, 19]. In 2013, Le et al. [20] applied the Hi-C technique to obtain interaction maps and models of *C. crescentus* with a resolution of 10kb and discovered for the first time the structural domains in bacteria similar to eukaryotic TADs, named CIDs (chromosomal interaction domains). In the same year, Cagliero et al. [21] obtained the first interaction map of *Escherichia coli* using 3C technology. In 2018, Lioy et al. [22] successfully obtained the high-quality interaction maps of *E. coli* with a resolution of 5kb, and reconstructed the spatial model using ShRec3D software. There are also several studies on the chromosome organization of *Bacillus subtilis*. In 2015, Marbouty et al. [23] used the Hi-C technique to study *B. subtilis*, and constructed a chromosome structure model using ShRec3D. They compared the structural model with super-resolution microscope data to reveal the folding patterns of chromosomes during replication initiation, chromosome organization, and DNA separation. Subsequently, Wang et al. conducted analysis on many interaction maps from *B. subtilis* cells, and uncovered the role of the SMC complex in chromosome compaction and replication processes [24, 25]. In a recent study, it was discovered that the Rok protein in *B. subtilis* forms anchored chromosomal loops at Mb scale, physically segregating large chromosomal regions [26]. With the emergence of an increasing number of bacterial 3C/Hi-C datasets [27-31], it is highly desired to have a recommendation on the 3D structure reconstruction tools to facilitate the prokaryotic 3D genomics. In this work, we compare some currently available software tools based on the same criteria.

## Current chromosome structure modeling methods and tools

### Constraint-based modeling

The constraints used in chromosome modeling are mainly derived from the DNA contact maps obtained through 3C/Hi-C experiments and the specific locus information from fluorescence microscopy. Among them, Hi-C data encompass genome-wide capture of chromosome contact probabilities, and thus have the characteristics of high throughput and large scale. However, their spatial interpretation is relatively intricate. Conversely, fluorescence locus information, such as fluorescence *in situ* hybridization (FISH) data, allows for the estimation of spatial distances between one or more pairs of distal loci, thereby explicitly indicating the physical proximity within genomic regions. Over the years, various algorithms have been proposed for inferring chromosome 3D structures from Hi-C contact data (**Figure 1, 2**). Below is a brief overview.

#### Multidimensional scaling

One common chromosome modeling approach is to convert DNA interaction frequencies into their spatial distances as constraints and then iteratively optimize the locus coordinates to find a conformation that satisfy these constraints. Using a distance matrix, multidimensional scaling (MDS) analysis can be performed. MDS is a multivariate statistical technique designed to map data from a high-dimensional space to a lower-dimensional space while preserving the distance relationships between original data points as much as possible. Depending on the nature of the optimization objective, MDS can be classified into two categories: metric MDS and non-metric MDS [32, 33].

The objective of metric MDS is to find a configuration of points in a multidimensional space [34]. It allows inferring the coordinates of points based on the given pairwise Euclidean distances between points. In the process of inferring 3D structures of chromosomes using metric MDS, the initial step involves assigning an expected spatial distance to each pair of DNA fragments (beads). This expected distance is derived from the DNA interaction matrix. Subsequently, all beads are positioned within 3D space to minimize the objective function, which is to ensure that the Euclidean distance between each pair of beads is as close as possible to the desired spatial distance. For example, Tanizawa et al. [35] used this method to reconstruct the chromosome model of fission yeast and included various restrictions, such as the minimum and maximum distances between adjacent beads [36], the minimum distance between an arbitrary pair of beads [37], and specific restrictions involving centromere, telomere, and the localization of ribosomal RNA coding region. However, the disadvantage of this method is that the function optimization results are dominated by pairs of beads with larger desired distances (smaller contact numbers, which are less reliable than larger contact numbers in experiments). To overcome this issue, Varoquaux et al. [38] proposed a variant that weights the contribution of a pair of beads (*i, j*) to the objective function inversely proportional to the squared distance between the corresponding beads. Additionally, ChromSDE proposed an optimization method based on metric MDS that maximizes the distances between pairs of beads without any interaction frequency data, by adding a regularization term to the weighted objective function [39]. Shavit et al. [40] developed an R package, FisHiCal, which integrates Hi-C and FISH data to perform FISH-based iterative Hi-C calibration. Next, the calibrated Hi-C data is used in the form of distances as input for the stress minimization function. Stress minimization is a common optimization technique for metric MDS, where it infers the 3D structure by adjusting the positions of data points in a lower-dimensional space.

Non-metric MDS (NMDS) provides another method for chromosome structure reconstruction. This method only uses the relative ranking information of data points in the distance matrix to construct a low-dimensional representation. For example, Ben-Elazar et al. [41] proposed a structural prediction method based on the assumption that if the observed contact frequency between a pair of beads *i* and *j* is higher than that between a pair of beads *k* and *v*, then the pair (*i, j*) should be closer in 3D space than the pair (*k, v*). Varoquaux et al. [38] proposed an optimization method to solve the NMDS problem by minimizing the Shepard-Kruskal scale cost function.

#### Poisson model and Bayesian inference

Hi-C data is known to exhibit certain systematic biases, including variations in the efficiency of enzyme digestion, GC content, and sequence uniqueness at the fragment ends [42]. Taking these biases into account, Hu et al. [43] proposed a probabilistic model in the BACH algorithm, which converts the structural inference problem into a maximum likelihood problem. This model uses a “beads-on-a-string” approach commonly used in chemistry and models the spatial distances (or contact frequencies) between beads as independent random variables following a Poisson distribution to correct known systematic biases. Similarly, Varoquaux et al. [38] adopted a Poisson model like that in BACH’s algorithm to estimate the maximum likelihood of the consensus structure directly. Carstens et al. [44] introduced a Bayesian probabilistic method that leverages the posterior probability distribution over the space of chromosome conformations and model parameters. This method combines information from single-cell Hi-C contacts and FISH measurements, providing a better definition of chromosome conformation.

However, high-resolution data often suffer from the existence of excess 0 contact counts in the contact matrix due to the sparsity of remote contacts between genomic loci [14]. Since the Poisson distribution only conforms to non-zero frequency, and if the proportion of zeros in the matrix is much higher than the theoretical probability of the Poisson distribution, the Poisson model may no longer be suitable for the data. To address this issue, Park et al. [45] proposed a tREX (truncated Random effect EXpression) model. This model uses truncated distribution to accommodate zeros in high-resolution data and adds a random effect component for counting. Consequently, the model exhibits strong robustness across different data resolutions.

#### Markov Chain Monte Carlo sampling

Markov Chain Monte Carlo (MCMC) methods have found broad application in computational biology such as RNA [46, 47] and protein [48, 49] structure prediction, phylogenetic inference [50, 51], and sequence alignment [52, 53]. MCMC is commonly used to generate robust candidate sets of structures based on noisy distance data. For example, Park et al. [45] used Hamiltonian MCMC when sampling the posterior value of the 3D structure coordinate set in the tREX model. Additionally, the BACH algorithm assumes that local genomic regions of interest (namely, TADs) exhibit consistent 3D chromosome structures in cell populations, and uses MCMC to infer potentially consistent 3D chromosome structure [43].

Rousseau et al. [54] developed a probabilistic model based on MCMC for linking Hi-C data to spatial distances, named as MCMC5C. Unlike optimization-based methods, MCMC5C models the uncertainty of spatial distance between two loci by assuming that the number of reads across two loci follows a Gaussian distribution. MCMC5C generates collections of different structures so that subclasses of structures can be discovered, and structural attributes and their distributions can be estimated to focus on statistically reasonable differences between attributes or data sets. However, estimating the Gaussian variance of each read count is challenging for MCMC5C, because a single Hi-C contact matrix does not provide sufficient information. Furthermore, the Gaussian model in MCMC5C is derived from the common 3D chromosome structure and cannot be used to measure structural changes in chromatin.

#### Lorentz equation

Hi-C data is generated from millions of cells in a single cell line, which can result in variations in genome structure and inconsistencies in chromosome contact. Consequently, it is challenging to satisfy the limitations of chromosome contact and its corresponding distance in a single structure. To address this issue, Trieu et al. [55] used the Lorentz function to design an objective function, which is more robust to outliers compared to the square error function. The Lorentzian function rewards the satisfaction of consistent constraints, and meanwhile ensures that the optimization process is not overly affected by inconsistent restraints. Additionally, this function is continuous and differentiable, which allows for the use of gradient-based optimization techniques effectively. Its scalability and noise-resisting feature make it a suitable approach for constructing whole 3D genome structures that involve noisy chromosome contacts.

#### Gradient descent/ascent

Gradient descent/ascent is a widely used iterative optimization algorithm in machine learning that aims to minimize/maximize the objective function or converge to its minimum/maximum value. In the 5C3D program [56], this method is used to find the best conformation of a virtual 3D DNA strand by minimizing the mismatch rate between the expected value in the distance matrix and the actual paired Euclidean distance. The MOGEN algorithm employs a contact-based optimization method that utilizes an adaptive step-size gradient ascent approach to continuously optimize the initial structure [57]. This algorithm does not require conversion of interaction frequency into distance before structure building, but rather tries to keep the distance between two contact regions below a threshold, which enables MOGEN to resist noise in the data. In the LorDG algorithm, gradient ascent optimization with an adaptive step size is used to adjust the position (namely, the *x, y, z* coordinates) of each bead to maximize a Lorentz objective function [55]. The 3DMax program employs an adaptive gradient ascent algorithm called AdaGrad, which adapts the learning rate automatically to each target parameter [58].

#### Other optimization algorithms

In addition to the methods mentioned above, there are several alternative approaches available for the reconstruction of chromosome 3D structures. Chromosome3D utilizes a distance geometry simulated annealing (DGSA) algorithm to reconstruct chromosome 3D structures from the chromosomal desired distance based on their interaction frequency [59]. It uses Hi-C distances as constraints for the simulated annealing (SA) optimization pipeline. The algorithm then optimizes the structural energy to obtain the expected 3D structure. Zhu et al. [60] introduced a modeling approach based on conformational energy and manifold learning. This framework interprets the spatial organization of chromosomes as the geometric structure of manifolds in 3D Euclidean space. It achieves this by converting Hi-C interaction frequencies into neighboring affinities of gene loci and then mapping them to Euclidean space to obtain the final 3D chromosome structure. Later on, Abbas et al. systematically integrated Hi-C and FISH data and proposed the GEM-FISH algorithm [61]. Due to the complementary nature of Hi-C and FISH data as constraints, this algorithm improved modeling result and revealed finer details of chromosomal packing.

### Thermodynamics-based modeling

The thermodynamics-based modeling approach considers chromosomes as polymer chains formed by the interactions of hundreds to thousands of molecular monomers. Therefore, it is essential to account for the steric configuration of molecular structures, including bond length, bond angle, and volume exclusion between non-adjacent molecules. Simultaneously, the conformational energy of polymers depends on the types of interactions, which consequently influence the stability, folding states, and overall structure of the polymer chain. Based on the polymer type or scale, thermodynamics-based modeling approaches can be categorized as follows (**Figure 3**).

**Figure 3.**
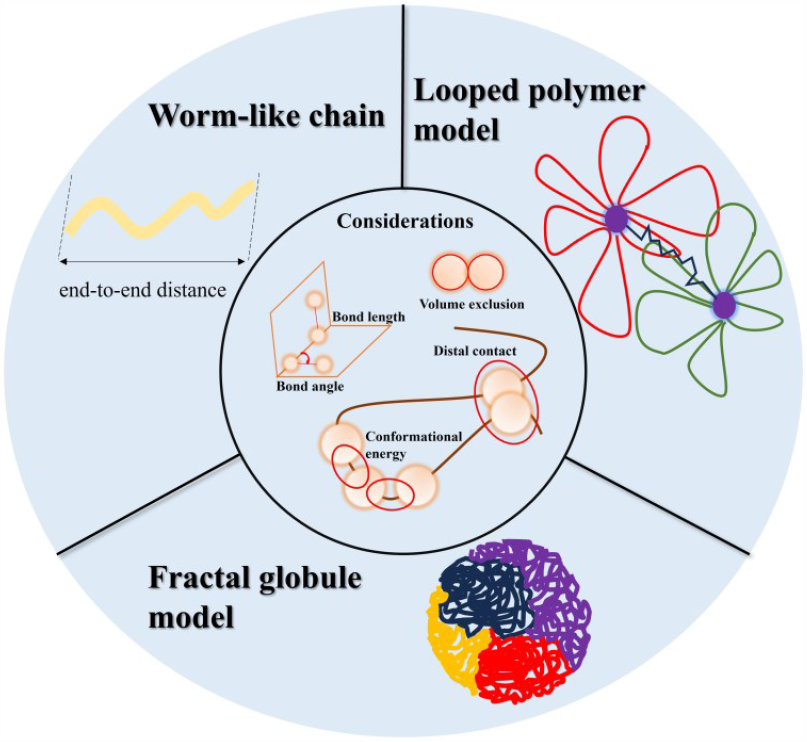
Representation of chromosomal polymer models in thermodynamic modeling. The outer layer provides an intuitive depiction of three model types, while the inner layer illustrates the factors considered during modeling.

#### Worm-like chain model

The Worm-like chain (WLC) model is a commonly used physical model for describing the conformation of semi-flexible polymers. This model assumes that each small segment of the polymer chain can be considered as a bent “worm-like” segment, connected by rigid joints. The primary concept behind the WLC model is to incorporate both the rigidity and flexibility in the chain description. Junier et al. [62] employed this model in combination with experimentally determined constraint parameters to perform numerical simulations of the chromosome polymer in *E. coli*. The constructed WLC model provides a coarse-grained description of protein-enveloped DNA that is used to observe chromosomal organization from a thermodynamic perspective during specific times of the cell cycle. By comparing with *in vivo* imaging data, it was revealed that the replication initiation and termination positions within structured macrodomains play a role in controlling the chromosome’s conformation and segregation.

In a research conducted by Buenemann and Lenz [63], DNA is also regarded as a semi-flexible polymer confined within the cell volume. In this context, a Monte Carlo method was developed for model testing, confirming a strong correlation between the position of gene on the chromosome and its spatial location within the cell volume. It is worth noting that while the WLC model is highly useful in capturing certain aspects of DNA mechanics, it simplifies some complexities of DNA behavior, such as the details involving base pairing and interactions with other molecules. In intricate scenarios or when specific interactions considered, more detailed models may be necessary.

#### Looped polymer model

To provide a better understanding of the compact folding of chromosomes, researchers have considered the possibility of introducing physical connections referred to as cis-interactions, which transform the linear arrangement of chromosomes into a looping structure [12]. The behavior of loops within polymer chains can be mathematically represented using various techniques, including topological parameters and statistical mechanics. The looped polymer model takes into account factors such as the length of loops, the probability of loop formation, and the impact of loops on the overall properties of the polymer. For instance, Sachs et al. [64] constructed a random-walk/giant-loop (RW/GL) model for human interphase chromosomes based on FISH data. In this model, chromatin fibers are conceptualized as undergoing random walks, and molecular motors moving along the chromatin fiber drive distal DNA regions together, resulting in the formation of large-scale chromatin loops. Subsequent analysis of FISH data led to the extension of the RW/GL model, giving rise to the multi-loop/subcompartment (MLS) model [65]. The MLS model posits that chromatin organization occurs at multiple hierarchical levels. It describes the genome as a series of looped structures, where contiguous loops form subcompartments. This hierarchical arrangement allows for efficient packaging of genetic material. Furthermore, the observed chromosome arms and subcompartments in the model align well with experimental results, indicating that polymer models are suitable for studying the 3D organization of the human interphase genome.

#### Fractal globule model

The fractal globule model is another theoretical framework for describing chromatin organization within the cell nucleus. This model, initially referred to as the crumpled globule, was proposed by Grosberg et al. [66], who argued that DNA confined within the cell nucleus needs an unknotted spatial structure to function effectively in a biological context. In contrast, the complex knotted structure of an equilibrium globule is deemed inadequate for maintaining the natural state of a functional biopolymer, as entanglements significantly reduce its reactivity to biochemical stress [67]. The fractal globule model suggests that chromatin organization does not exist in thermodynamic equilibrium. Instead, it maintains equilibrium through active processes, such as the active squeezing of chromatin loops by protein complexes. Later, with the application of Hi-C techniques, Lieberman-Aiden et al. [5] discovered that this fractal state indeed aligned with Hi-C data obtained from human cells. The fractal structure of the model implies that regions of the DNA chain that are close in 1D distance also tend to occupy adjacent regions in 3D space, enhancing accessibility to specific genomic regions within the cell nucleus, which is crucial for gene regulation and other cellular processes.

In prokaryotes, a nucleotide-resolution model of the *E. coli* chromosome, similar to the fractal globule, was described by Hacker et al. [68]. Due to the high complexity of the model structure, the degree of knotting could not be determined. However, in the *E. coli* model, the relationship function *P*(*s*) between the contact probability of two loci and their 1D distance along the genome satisfies the *s*^*-1*^ scaling law, consistent with the properties of the fractal globule model. By capturing the elastic topology and physical properties of double-stranded and supercoiled DNA, the multi-scale polymer model was developed. This model identified the four chromosome regions corresponding to macrodomains, and these regions automatically separated from each other and occupied specific positions within the nucleoid. Inspired by the *C. crescentus* model proposed by the Laub lab [20], Hacker et al. divided the chromosome into plectoneme-abundant and plectoneme-free regions (PFRs). PFRs (or the non-supercoiling regions) are identified based on RNAP binding site recognition from ChIP-chip data and considered corresponding to highly transcriptionally active regions. This nucleotide-resolution model represents a pioneering investigation of the physical properties of a bacterial chromosome at the nucleotide level, discussing the characteristics of a chromosome at both macroscopic and microscopic levels.

## Tests and evaluation

### List of evaluated software tools

The evaluated software tools are listed in **Table 1**. These tools mainly utilize Hi-C interaction matrices as the input data, and demonstrate robust reproducibility and ease of visualization in the generated structure models. These modeling algorithms are implemented with diverse programming languages, exemplifying certain representativeness within the field. Moreover, their open-source nature ensures accessibility to the source code. The software tools are all run under default parameters.

**Table 1.**
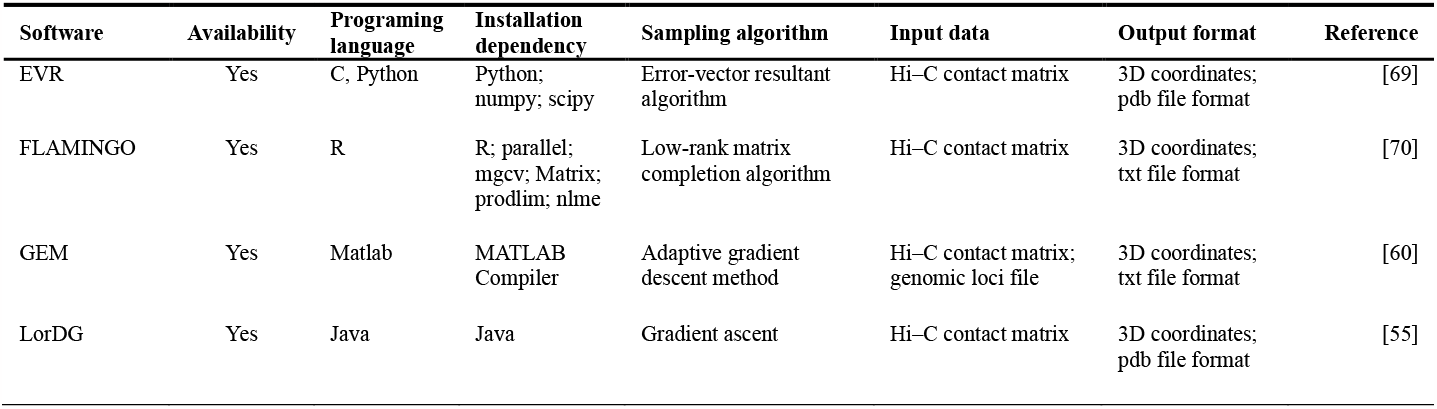
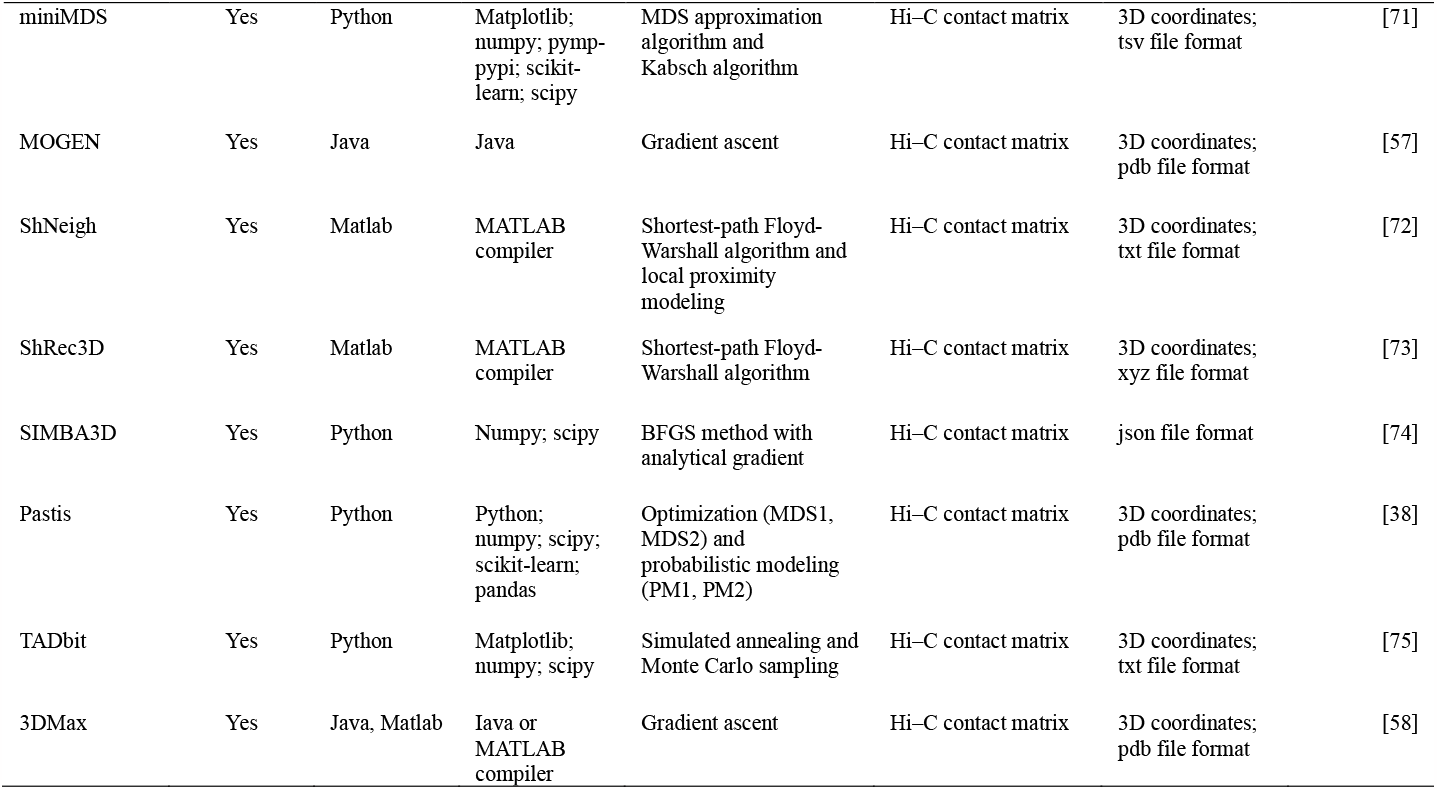
Information on the software tools for chromosome 3D structure reconstruction.

The installation instructions for the tested software and a list of untested software along with the reasons for not testing are presented in the **Supplementary information** (**Appendix 1** and **Appendix 2**). Furthermore, we have packaged the tested software and their installation environments into a Docker image which can be downloaded from the Docker Hub website.

## Data sets used in evaluation

The evaluation data consisted of three datasets: publicly available chromosomal interaction data for *E. coli* (GEO accession number: GSE107301) with a resolution of 5kb [22], *C. crescentus* (GEO accession number: GSE45966) with a resolution of 10kb [20] and a simulating dataset of spiral ring structure [69].

### Platform and measures

In this study, we aim to evaluate the accuracy and robustness of several chromosome 3D structure reconstruction methods. Our evaluation process is underpinned by the utilization of two distinct criteria: the Pearson Correlation Coefficient (PCC) and the Root Mean Square Deviation (RMSD) value. To assess accuracy, we compared the chromosome 3D structure of *E. coli* and *C. crescentus* generated by each software tool with corresponding published fluorescence microscopy experimental data [76, 77]. We mapped the fluorescence labeled sites onto the reconstructed structures to determine the 3D coordinates of these sites, and then calculated the PCC between the spatial distance in structure and the experimental distance measured by fluorescence microscopy. To assess robustness, following a previous work [69], we added different levels of noise ranging from -0.5 × P × IF_max_ to 0.5 × P × IF_max_ (P is noise level and IF_max_ is the maximum value in IF matrix) to the IF matrix generated from the structure of spiral ring, and then used the noisy data for 3D structure reconstruction using the software tools described above. We compared the reconstructed structure from noisy data with the original structure. At each noise level, we generated 100 sets of data for structural reconstruction and calculated the average RMSD value.

We also evaluated the running speed of the software tools, which is an important factor affecting user experience. We generated the IF matrix of spiral ring ranging from 100-2000 bins and used the software tools to reconstruct the 3D structure. For each bin number, we generated 10 groups of data and compared the running time on the same platform. The platform used was an Ubuntu18.04 (64-bit) system with Intel(R) Core (TM) i5-2400 CPU at 3.10GHz and 20 GB DDR3 1600MH memory, and an Nvidia GeForce GTX 1050Ti GPU.

### Accuracy

The accuracy of each modeling software was assessed by comparing the reconstructed 3D structures of the two bacterial chromosomes with corresponding fluorescence microscopy data (**Table 2**). The EVR algorithm, designed specifically for prokaryotic chromosome reconstruction, exhibits the highest correlation between modeling and experimental distances in *E. coli* and the fourth highest in *C. crescentus*. GEM, LorDG and two statistical methods using Poisson distribution (Pastis_PM1 and PM2) also perform well. Although PM2 can automatically adjust the formula to infer the best model compared with PM1, our test reveals small disparity between PM1 and PM2. Compared with the performance in *E. coli*, Pastis_MDS and Pastis_NMDS exhibits inferior correlation when applied to *C. crescentus*. This discrepancy may likely be attributed to the instability of these algorithms in processing data with different resolutions. Among the other algorithms evaluated, FLAMINGO, miniMDS, ShNeigh, SIMBA3D, TADbit and 3DMax also showed moderate and significant correlations, despite not being specifically developed for prokaryotes. MOGEN and ShRec3D displayed a relatively low correlation, likely due to its unsuitability for prokaryotes. Moreover, the default parameter configuration of some algorithms may be suboptimal for bacterial chromosome modeling, which may also contribute to their weaker correlations.

**Table 2.** Evaluation of modeling accuracy based on fluorescence microscopy data.

| Algorithm | *E. coli* |  | *C. crescentus* |  |
| --- | --- | --- | --- | --- |
|  | *PCC* | *p-value* | *PCC* | *p-value* |
| EVR | 0.86936 | 1.08E-10 | 0.91633 | 3.80E-43 |
| FLAMINGO | 0.76219 | 4.00E-07 | 0.80426 | 1.02E-19 |
| GEM | 0.82388 | 1.12E-07 | 0.88683 | 6.24E-25 |
| LorDG | 0.82951 | 3.18E-06 | 0.93491 | 5.67E-46 |
| miniMDS | 0.79629 | 5.16E-06 | 0.81751 | 7.52E-25 |
| MOGEN | 0.61573 | 2.46E-02 | 0.42921 | 2.33E-06 |
| ShNeigh1 | 0.77635 | 1.77E-07 | 0.69017 | 2.64E-07 |
| ShNeigh2 | 0.83073 | 4.00E-09 | 0.7969 | 1.00E-14 |
| ShRec3D | 0.6026 | 6.81E-03 | 0.54118 | 6.33E-04 |
| SIMBA3D | 0.71325 | 5.50E-04 | 0.69153 | 8.57E-06 |
| Pastis_MDS | 0.82801 | 8.38E-08 | 0.60478 | 4.88E-06 |
| Pastis_NMDS | 0.78343 | 7.07E-06 | 0.63657 | 1.54E-07 |
| Pastis_PM1 | 0.85078 | 2.05E-08 | 0.91783 | 7.88E-43 |
| Pastis_PM2 | 0.84656 | 3.99E-08 | 0.91816 | 2.11E-42 |
| TADbit | 0.73948 | 1.01E-03 | 0.78287 | 5.14E-15 |
| 3DMax | 0.82533 | 9.11E-08 | 0.88724 | 2.27E-31 |

### Robustness

To evaluate the robustness of various chromosome 3D structure modeling tools, we generated datasets of spiral ring with varying levels of noise. As shown in **Figure 4**, without adding noise, the NMDS algorithm of the pastis software tool (Pastis_NMDS) performs the best, followed by the EVR algorithm, when compared with the original standard structure. Furthermore, the algorithms of 3DMax, ShRec3D, SIMBA3D, miniMDS, and Pastis_PM2 all perform well, and the RMSD values of structures produced by the MOGEN, TADbit, LorDG, ShNeigh2, GEM, FLAMINGO algorithms are also at a low level. However, the ShNeigh1, Pastis_PM1, Pastis_MDS algorithms produce structures with relatively high RMSD values, indicating significant differences from the original structure.

**Figure 4.**
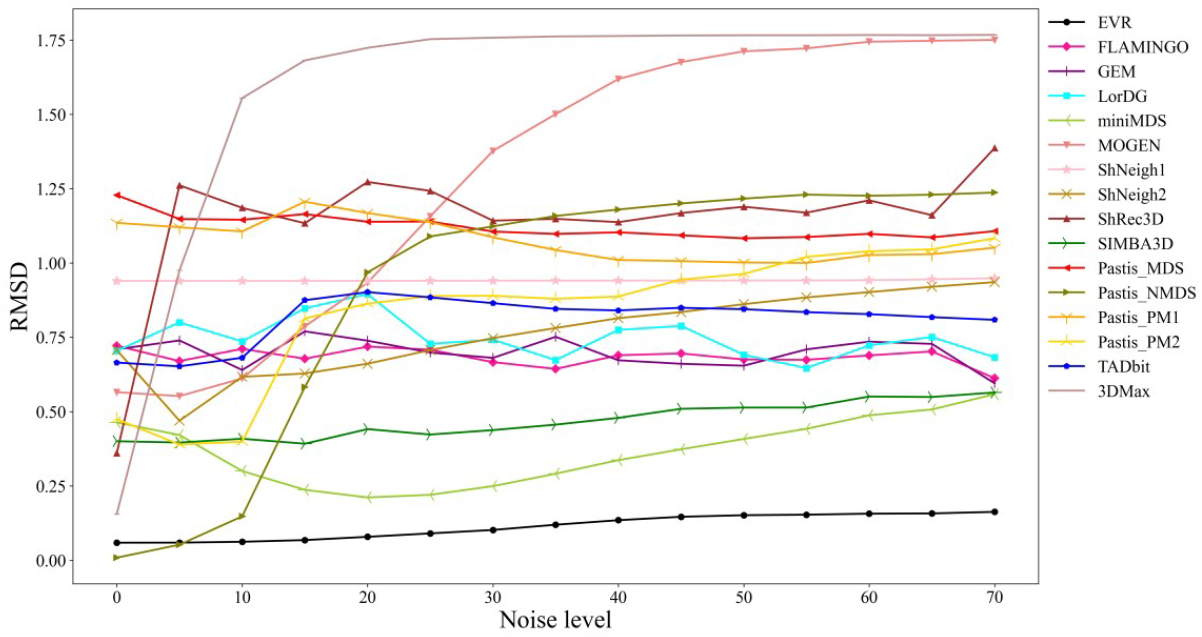
Comparison between the reconstructed structure (from noisy data) and the original structure using RMSD. Both the absolute RMSD value and its trend with the increase of noise level should be considered.

After adding different levels of noise to the dataset, the EVR algorithm shows the best robustness and lowest RMSD level. The algorithms of miniMDS, SIMBA3D, FLAMINGO, GEM, LorDG, ShNeigh, TADbit, Pastis_MDS, and Pastis_ PM1 also show good robustness. However, the ShRec3D, Pastis_NMDS, Pastis_PM2, MOGEN, 3DMax algorithms demonstrate poor robustness in generating structures under different levels of noise.

### Speed

We used various software/algorithms to reconstruct the 3D structure of standard spiral ring data with varying numbers of bins, and the resulting time efficiency are presented in **Figure 5**. The results show that the EVR algorithm is the fastest among all the tested algorithms. Additionally, the running time of EVR is not sensitive to the number of bins, within a reasonable range. The computational speed of miniMDS, Mogen, ShNeigh1, ShRec3D, 3DMax, and FLAMINGO algorithms is slower than that of the EVR algorithm but still relatively fast. In contrast, the Pastis_MDS, LorDG, TADbit, Pastis_NMDS, SIMBA3D and Pastis_PM1 algorithms run slowly. The Pastis_PM2, ShNeigh2, and GEM algorithms are the slowest among all the tested algorithms. Moreover, the runtime of these algorithms increases significantly as the number of bins increases.

**Figure 5.**
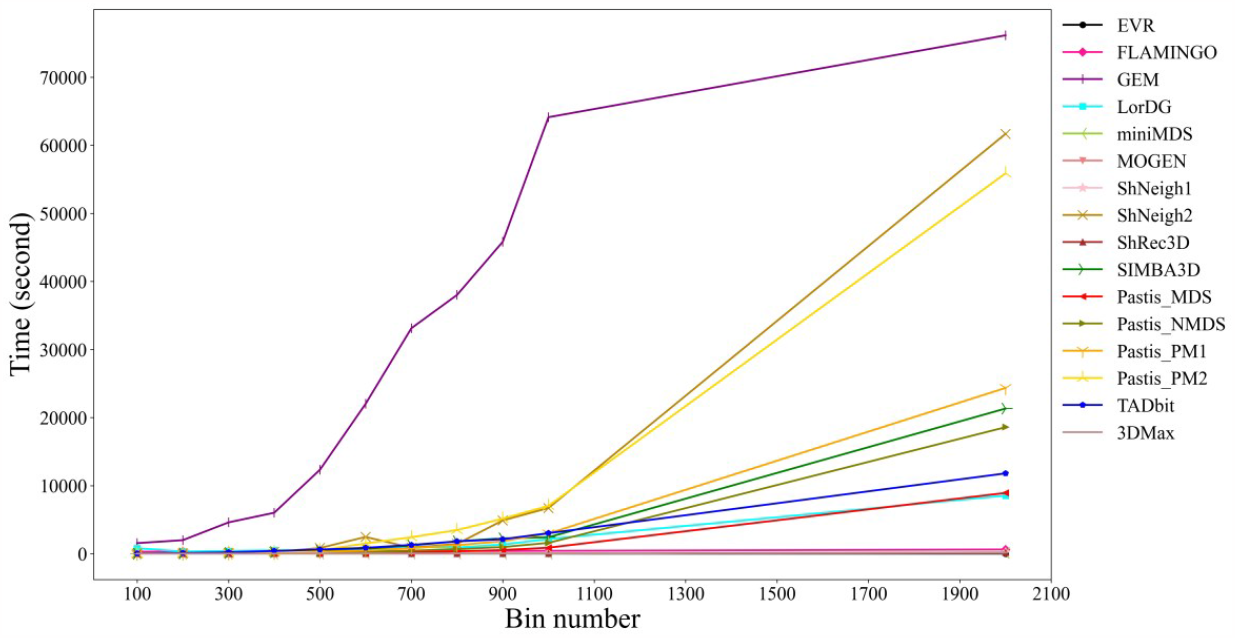
The time required to reconstruct the 3D model of the standard spiral ring structure with various bin numbers. This figure shows the time efficiency of different modeling algorithms.

### Limitations and problems

This work provides an overview of the current state of mainstream chromosome 3D structure reconstruction software, which are predominantly written in Matlab, Python, R, Perl, Java, C++ and other languages. However, the installation and use of these software pose some challenges. Firstly, many of the software download links are invalid or the program packages cannot be found. Secondly, some software requires complex dependencies that are difficult to resolve, or may encounter unknown errors when run after installation. Thirdly, certain software may require significant parameter tuning to identify optimal parameters for specific data sets, making the exploration process time-consuming and challenging. Additionally, the input and output data formats vary largely across different software tools, making it difficult to compare the modeling results.

With regards to input file format, the Hi-C interaction data are usually provided in text files with matrix or list formats. Some software requires additional data such as restriction enzyme cutting frequency, GC content, sequence uniqueness, FISH data and Hi-C contact matrix files containing identified TADs. In prokaryotic chromosome structure modeling, some relevant data may be missing. For example, Chrom3D requires lamin-associated domain (LAD) information, making it unsuitable for modeling prokaryotic chromosomes [78].

In terms of output file format, all current modeling software generates structures containing spatial coordinate information of bins, which are usually saved in simple formats similar to PDB files, such as the modified PDB format, “xyz” format, “mat” format and “txt” format. While some of these formats can be visualized directly through tools such as PyMol and Chimera, others such as the txt format require additional processing and conversion before visualization. This increases the difficulty of work involved in comparing and analyzing results.

### Perspective

#### Modeling bacteria chromosome with replication forks

Bacterial chromosome replication typically originates at a single point and proceeds bidirectionally until the two replication forks converge in the terminal region [79]. Following replication initiation, active mechanisms rapidly separate the two copies [80, 81]. In eukaryotes, experiments have been conducted to distinguish between sister chromatids. As sister chromatids share the same base sequence during replication, it can be challenging to differentiate the chromosomal interactions they are involved in. To address this issue, researchers have proposed the sister-chromatid-sensitive Hi-C (scsHi-C) [82] and SisterC [83] methods. The former introduces a sister-chromatid-specific marker and culture cells in the presence of DNA nucleotide analogs. A round of DNA replication is performed to label the Watson and Crick strands in the two sister chromatids. The latter utilizes a combination of Hi-C with 5-bromo-2’-deoxyuridine (BrdU)-incorporated DNA and Hoechst/UV treatment to distinguish the interactions between sister chromatids (inter-sister interactions) and along individual sister chromatids (intra-sister interactions). Espinosa et al. proposed a high-throughput method for monitoring sister chromatid contact (Hi-SC2) [84]. Using a multi-chromosome species *Vibrio cholerae* as a model, they monitored local variations in sister chromatid cohesion at high resolution throughout the genome. In terms of modeling, Wasim et al. [85] clarified the multi-scale organization of *E. coli* chromosomes at different replication stages by integrating the beads-on-a-spring model and the Hi-C interaction matrix. However, no software has been developed yet to construct a chromosome model in replication directly from Hi-C interaction data.

### Integration of structure models with multi-omics data

With the advancement of technologies such as 3C and Hi-C, there has been increasing attention to the impact of the unique 3D structure of bacterial chromosomes on their metabolic activities. Recently, modeling algorithms have been used to reconstruct bacterial chromosome 3D structures, resulting in successful construction of more such structures [15, 20, 27, 29]. Integration of chromosome 3D structure data with other data has become a new research trend. Several studies have already combined chromosome sequencing data with other data sources. For instance, Xie et al. [86] integrated the genome conformation capture data of *E. coli* with its genome, biological pathway, and protein interaction data, leading to the discovery of the spatial characteristics of *E. coli* genome organization. Meanwhile, Hołówka and Płachetka [87] used molecular biology techniques along with high-throughput DNA sequencing methods to analyze bacterial chromosome structures in a precise manner at both local and global scales. Tian et al. [88] investigated the spatial organization characteristics of bacterial transcriptional regulatory network (TRN) using gene regulation and chromatin interaction data. Under different physiological conditions, the spatial organization features of bacterial TRNs remain relatively stable. The research results provide new insights into the connection between transcriptional regulation and chromosome spatial organization in bacteria.

### Dynamics of chromosome structure

Chromosome organization and dynamics have primarily been studied in eukaryotes due to their observable aggregation and separation by microscopy. In eukaryotic cells, 3C techniques have been used to analyze the dynamics of high-order chromosome structures, resulting in significant progress. For instance, Nagano et al. [89] utilized high-throughput single-cell Hi-C to generate and analyze single-cell contact diagrams of thousands of cells, revealing high genome folding heterogeneity at the single-cell level. The study confirmed that the high heterogeneity is a product of deterministic dynamics and random effects and that the cell cycle has the greatest influence on the deterministic dynamics of embryonic stem cells in mice. Recent Hi-C studies have also demonstrated that enhancer-promoter interactions and gene expression are dynamically regulated by high-order chromosome topologies [90].

Initially, bacterial chromosomes were thought to be compact and unstructured due to methodological limitations. However, with the advent of living cell fluorescence microscopy [91], it was discovered that bacterial chromosomes are dynamic and highly organized [92]. Current research aims to understand the molecular mechanisms behind bacterial chromosome structuring and its dynamics [93]. Recent advances in 3C, Hi-C, and single-cell Hi-C technology have provided new possibility for the study of bacterial chromosome dynamics. For example, the structure dynamics of bacterial chromosomes during the cell cycle based on reconstructed 3D models is still a relatively unexplored area. Future research should focus on this direction.

## Supporting information

Appendix 1 - Software Installation Instructions.pdf

Appendix 2 - Untested Software.pdf

## Acknowledgements

This work was supported by the National Natural Science Foundation of China (Grant 31971184). The funders had no role in study design, data collection and interpretation, or the decision to submit the work for publication.

## References

[1] Ji X, Dadon DB, Powell BE, et al. 3D chromosome regulatory landscape of human pluripotent cells. Cell Stem Cell 2016;18(2):262–75.

[2] Dekker J, Rippe K, Dekker M, et al. Capturing chromosome conformation. Science 2002;295(5558):1306–11.

[3] Simonis M, Klous P, Splinter E, et al. Nuclear organization of active and inactive chromatin domains uncovered by chromosome conformation capture-on-chip (4C). Nat Genet 2006;38(11):1348–54.

[4] Dostie J, Richmond TA, Arnaout RA, et al. Chromosome Conformation Capture Carbon Copy (5C): a massively parallel solution for mapping interactions between genomic elements. Genome Res 2006;16(10):1299–309.

[5] Lieberman-Aiden E, van Berkum NL, Williams L, et al. Comprehensive mapping of long-range interactions reveals folding principles of the human genome. Science 2009;326(5950):289–93.

[6] Lin D, Hong P, Zhang S, et al. Digestion-ligation-only Hi-C is an efficient and cost-effective method for chromosome conformation capture. Nat Genet 2018;50(5):754–63.

[7] Dixon JR, Selvaraj S, Yue F, et al. Topological domains in mammalian genomes identified by analysis of chromatin interactions. Nature 2012;485(7398):376–80.

[8] Rocha PP, Raviram R, Bonneau R, et al. Breaking TADs: insights into hierarchical genome organization. Epigenomics 2015;7(4):523–6.

[9] Dixon JR, Gorkin DU, Ren B. Chromatin domains: the unit of chromosome organization. Mol Cell 2016;62(5):668–80.

[10] Gonzalez-Sandoval A, Gasser SM. On TADs and LADs: spatial control over gene expression. Trends Genet 2016;32(8):485–95.

[11] Serra F, Di Stefano M, Spill YG, et al. Restraint-based three-dimensional modeling of genomes and genomic domains. FEBS Lett 2015;589(20, Part A):2987–95.

[12] Rosa A, Zimmer C. Computational models of large-scale genome architecture. Int Rev Cell Mol Biol 2014;307:275–349.

[13] Bianco S, Chiariello AM, Annunziatella C, et al. Predicting chromatin architecture from models of polymer physics. Chromosome Res 2017;25(1):1–10.

[14] Oluwadare O, Highsmith M, Cheng J. An overview of methods for reconstructing 3-D chromosome and genome structures from Hi-C data. Biol Proced Online 2019;21:7.

[15] Yildirim A, Feig M. High-resolution 3D models of Caulobacter crescentus chromosome reveal genome structural variability and organization. Nucleic Acids Res 2018;46(8):3937–52.

[16] Bonev B, Mendelson Cohen N, Szabo Q, et al. Multiscale 3D genome rewiring during mouse neural development. Cell 2017;171(3):557–72.

[17] Dong Q, Li N, Li X, et al. Genome-wide Hi-C analysis reveals extensive hierarchical chromatin interactions in rice. Plant J 2018;94(6):1141–56.

[18] Umbarger MA, Toro E, Wright MA, et al. The three-dimensional architecture of a bacterial genome and its alteration by genetic perturbation. Mol Cell 2011;44(2):252–64.

[19] Russel D, Lasker K, Webb B, et al. Putting the pieces together: integrative modeling platform software for structure determination of macromolecular assemblies. PLoS Biol 2012;10(1):e1001244.

[20] Le TB, Imakaev MV, Mirny LA, et al. High-resolution mapping of the spatial organization of a bacterial chromosome. Science 2013;342(6159):731–4.

[21] Cagliero C, Grand RS, Jones MB, et al. Genome conformation capture reveals that the Escherichia coli chromosome is organized by replication and transcription. Nucleic Acids Res 2013;41(12):6058–71.

[22] Lioy VS, Cournac A, Marbouty M, et al. Multiscale structuring of the E. coli chromosome by nucleoid-associated and condensin proteins. Cell 2018;172(4):771–83.

[23] Marbouty M, Cattoni DI, Cournac A, et al. Condensin- and replication-mediated bacterial chromosome folding and origin condensation revealed by Hi-C and super-resolution imaging. Mol Cell 2015;59(4):588–602.

[24] Wang X, Le TB, Lajoie BR, et al. Condensin promotes the juxtaposition of DNA flanking its loading site in Bacillus subtilis. Genes Dev 2015;29(15):1661–75.

[25] Wang X, Brandão HB, Le TBK, et al. Bacillus subtilis SMC complexes juxtapose chromosome arms as they travel from origin to terminus. Science 2017;355(6324):524–7.

[26] Dugar G, Hofmann A, Heermann DW, et al. A chromosomal loop anchor mediates bacterial genome organization. Nat Genet 2022;54(2):194–201.

[27] Val ME, Marbouty M, de Lemos Martins F, et al. A checkpoint control orchestrates the replication of the two chromosomes of Vibrio cholerae. Sci Adv 2016;2(4):e1501914.

[28] Ren Z, Liao Q, Karaboja X, et al. Conformation and dynamic interactions of the multipartite genome in Agrobacterium tumefaciens. Proc Natl Acad Sci U S A 2022;119(6):e2115854119.

[29] Trussart M, Yus E, Martinez S, et al. Defined chromosome structure in the genome-reduced bacterium Mycoplasma pneumoniae. Nat Commun 2017;8:14665.

[30] Lioy VS, Junier I, Lagage V, et al. Distinct activities of bacterial condensins for chromosome management in Pseudomonas aeruginosa. Cell Rep 2020;33(5):108344.

[31] Conin B, Billault-Chaumartin I, El Sayyed H, et al. Extended sister-chromosome catenation leads to massive reorganization of the E. coli genome. Nucleic Acids Res 2022;50(5):2635–50.

[32] Shepard RN. The analysis of proximities: Multidimensional scaling with an unknown distance function. Part I. Psychometrika 1962;27(2):125–40.

[33] Kruskal JB. Multidimensional scaling by optimizing goodness of fit to a nonmetric hypothesis. Psychometrika 1964;29(1):1–27.

[34] Steyvers M. Multidimensional scaling. In: Nadel L. (ed) The Encyclopedia of Cognitive Science. Macmillan, 2002.

[35] Tanizawa H, Iwasaki O, Tanaka A, et al. Mapping of long-range associations throughout the fission yeast genome reveals global genome organization linked to transcriptional regulation. Nucleic Acids Res 2010;38(22):8164–77.

[36] Bystricky K, Heun P, Gehlen L, et al. Long-range compaction and flexibility of interphase chromatin in budding yeast analyzed by high-resolution imaging techniques. Proc Natl Acad Sci U S A 2004;101(47):16495–500.

[37] Dehghani H, Dellaire G, Bazett-Jones DP. Organization of chromatin in the interphase mammalian cell. Micron 2005;36(2):95–108.

[38] Varoquaux N, Ay F, Noble WS, et al. A statistical approach for inferring the 3D structure of the genome. Bioinformatics 2014;30(12):i26–33.

[39] Zhang Z, Li G, Toh K-C, et al. Inference of spatial organizations of chromosomes using semi-definite embedding approach and hi-c data. In: Proceedings of the 17th international conference on Research in Computational Molecular Biology. Beijing, China, 2013, p. 317–32. Springer-Verlag.

[40] Shavit Y, Hamey FK, Lio P. FisHiCal: an R package for iterative FISH-based calibration of Hi-C data. Bioinformatics 2014;30(21):3120–2.

[41] Ben-Elazar S, Yakhini Z, Yanai I. Spatial localization of co-regulated genes exceeds genomic gene clustering in the Saccharomyces cerevisiae genome. Nucleic Acids Res 2013;41(4):2191–201.

[42] Yaffe E, Tanay A. Probabilistic modeling of Hi-C contact maps eliminates systematic biases to characterize global chromosomal architecture. Nat Genet 2011;43(11):1059–65.

[43] Hu M, Deng K, Qin Z, et al. Bayesian inference of spatial organizations of chromosomes. PLoS Comput Biol 2013;9(1):e1002893.

[44] Carstens S, Nilges M, Habeck M. Inferential structure determination of chromosomes from single-cell Hi-C data. PLoS Comput Biol 2016;12(12):e1005292.

[45] Park J, Lin S. Impact of data resolution on three-dimensional structure inference methods. BMC Bioinform 2016;17(1):70.

[46] Meyer IM, Miklos I. SimulFold: simultaneously inferring RNA structures including pseudoknots, alignments, and trees using a Bayesian MCMC framework. PLoS Comput Biol 2007;3(8):e149.

[47] Metzler D, Nebel ME. Predicting RNA secondary structures with pseudoknots by MCMC sampling. J Math Biol 2008;56(1-2):161–81.

[48] Boomsma W, Mardia KV, Taylor CC, et al. A generative, probabilistic model of local protein structure. Proc Natl Acad Sci U S A 2008;105(26):8932–7.

[49] Robinson DM, Jones DT, Kishino H, et al. Protein evolution with dependence among codons due to tertiary structure. Mol Biol Evol 2003;20(10):1692–704.

[50] Huelsenbeck JP, Ronquist F, Nielsen R, et al. Bayesian inference of phylogeny and its impact on evolutionary biology. Science 2001;294(5550):2310–4.

[51] Rodrigue N, Kleinman CL, Philippe H, et al. Computational methods for evaluating phylogenetic models of coding sequence evolution with dependence between codons. Mol Biol Evol 2009;26(7):1663–76.

[52] Zhu J, Liu JS, Lawrence CE. Bayesian adaptive sequence alignment algorithms. Bioinformatics 1998;14(1):25–39.

[53] Lunter G, Miklós I, Drummond A, et al. Bayesian coestimation of phylogeny and sequence alignment. BMC Bioinform 2005;6:83.

[54] Rousseau M, Fraser J, Ferraiuolo MA, et al. Three-dimensional modeling of chromatin structure from interaction frequency data using Markov chain Monte Carlo sampling. BMC Bioinform 2011;12:414.

[55] Trieu T, Cheng J. 3D genome structure modeling by Lorentzian objective function. Nucleic Acids Res 2017;45(3):1049–58.

[56] Fraser J, Rousseau M, Shenker S, et al. Chromatin conformation signatures of cellular differentiation. Genome Biol 2009;10(4):R37.

[57] Trieu T, Cheng J. MOGEN: a tool for reconstructing 3D models of genomes from chromosomal conformation capturing data. Bioinformatics 2016;32(9):1286–92.

[58] Oluwadare O, Zhang Y, Cheng J. A maximum likelihood algorithm for reconstructing 3D structures of human chromosomes from chromosomal contact data. BMC Genom 2018;19(1):161.

[59] Adhikari B, Trieu T, Cheng J. Chromosome3D: reconstructing three-dimensional chromosomal structures from Hi-C interaction frequency data using distance geometry simulated annealing. BMC Genom 2016;17(1):886.

[60] Zhu G, Deng W, Hu H, et al. Reconstructing spatial organizations of chromosomes through manifold learning. Nucleic Acids Res 2018;46(8):e50–e.

[61] Abbas A, He X, Niu J, et al. Integrating Hi-C and FISH data for modeling of the 3D organization of chromosomes. Nat Commun 2019;10(1):2049.

[62] Junier I, Boccard F, Espeli O. Polymer modeling of the E. coli genome reveals the involvement of locus positioning and macrodomain structuring for the control of chromosome conformation and segregation. Nucleic Acids Res 2014;42(3):1461–73.

[63] Buenemann M, Lenz P. A geometrical model for DNA organization in bacteria. PLoS One 2010;5(11):e13806.

[64] Sachs RK, van den Engh G, Trask B, et al. A random-walk/giant-loop model for interphase chromosomes. Proc Natl Acad Sci U S A 1995;92(7):2710–4.

[65] Münkel C, Langowski J. Chromosome structure predicted by a polymer model. Phys Rev E 1998;57(5):5888–96.

[66] Grosberg AY, Nechaev SK, Shakhnovich EI. The role of topological constraints in the kinetics of collapse of macromolecules. J Phys 1988;49:2095–100.

[67] Grosberg AY, Rabin Y, Havlin S, et al. Crumpled globule model of the three-dimensional structure of DNA. EPL 1993;23(5):373–8.

[68] Hacker WC, Li S, Elcock AH. Features of genomic organization in a nucleotide-resolution molecular model of the Escherichia coli chromosome. Nucleic Acids Res 2017;45(13):7541–54.

[69] Hua KJ, Ma BG. EVR: reconstruction of bacterial chromosome 3D structure models using error-vector resultant algorithm. BMC Genom 2019;20(1):738.

[70] Wang H, Yang J, Zhang Y, et al. Reconstruct high-resolution 3D genome structures for diverse cell-types using FLAMINGO. Nat Commun 2022;13(1):2645.

[71] Rieber L, Mahony S. miniMDS: 3D structural inference from high-resolution Hi-C data. Bioinformatics 2017;33(14):i261–i6.

[72] Li F-Z, Liu Z-E, Li X-Y, et al. Chromatin 3D structure reconstruction with consideration of adjacency relationship among genomic loci. BMC Bioinform 2020;21(1):272.

[73] Lesne A, Riposo J, Roger P, et al. 3D genome reconstruction from chromosomal contacts. Nat Methods 2014;11(11):1141–3.

[74] Rosenthal M, Bryner D, Huffer F, et al. Bayesian estimation of three-dimensional chromosomal structure from single-cell Hi-C data. J Comput Biol 2019;26(11):1191–202.

[75] Serra F, Bau D, Goodstadt M, et al. Automatic analysis and 3D-modelling of Hi-C data using TADbit reveals structural features of the fly chromatin colors. PLoS Comput Biol 2017;13(7):e1005665.

[76] Espeli O, Mercier R, Boccard F. DNA dynamics vary according to macrodomain topography in the E. coli chromosome. Mol Microbiol 2008;68(6):1418–27.

[77] Viollier PH, Thanbichler M, McGrath PT, et al. Rapid and sequential movement of individual chromosomal loci to specific subcellular locations during bacterial DNA replication. Proc Natl Acad Sci U S A 2004;101(25):9257–62.

[78] Paulsen J, Sekelja M, Oldenburg AR, et al. Chrom3D: three-dimensional genome modeling from Hi-C and nuclear lamin-genome contacts. Genome Biol 2017;18(1):21.

[79] O’Donnell M. Replisome architecture and dynamics in Escherichia coli. J Biol Chem 2006;281(16):10653–6.

[80] Gordon GS, Sitnikov D, Webb CD, et al. Chromosome and low copy plasmid segregation in E. coli: visual evidence for distinct mechanisms. Cell 1997;90(6):1113–21.

[81] Webb CD, Graumann PL, Kahana JA, et al. Use of time-lapse microscopy to visualize rapid movement of the replication origin region of the chromosome during the cell cycle in Bacillus subtilis. Mol Microbiol 1998;28(5):883–92.

[82] Mitter M, Gasser C, Takacs Z, et al. Conformation of sister chromatids in the replicated human genome. Nature 2020;586(7827):139–44.

[83] Oomen ME, Hedger AK, Watts JK, et al. Detecting chromatin interactions between and along sister chromatids with SisterC. Nat Methods 2020;17(10):1002–9.

[84] Espinosa E, Paly E, Barre FX. High-resolution whole-genome analysis of sister-chromatid contacts. Mol Cell 2020;79(5):857–69.

[85] Wasim A, Gupta A, Mondal J. A Hi–C data-integrated model elucidates E. coli chromosome’s multiscale organization at various replication stages. Nucleic Acids Res 2021;49(6):3077–91.

[86] Xie T, Fu LY, Yang QY, et al. Spatial features for Escherichia coli genome organization. BMC Genom 2015;16(1):37.

[87] Hołówka J, Płachetka M. Structure of bacterial chromosome: An analysis of DNA-protein interactions in vivo. Postepy Hig Med Dosw (Online) 2017;71(0):1005–14.

[88] Tian L, Liu T, Hua KJ, et al. The spatial organization of bacterial transcriptional regulatory networks. Microorganisms 2022;10(12):2366.

[89] Nagano T, Lubling Y, Várnai C, et al. Cell-cycle dynamics of chromosomal organization at single-cell resolution. Nature 2017;547(7661):61–7.

[90] Yokoshi M, Fukaya T. Dynamics of transcriptional enhancers and chromosome topology in gene regulation. Dev Growth Differ 2019;61(5):343–52.

[91] Ettinger A, Wittmann T. Chapter 5 - Fluorescence live cell imaging. In: Waters J. C., Wittman T. eds. Methods Cell Biol. Academic Press, 2014, 77–94.

[92] Thanbichler M, Viollier PH, Shapiro L. The structure and function of the bacterial chromosome. Curr Opin Genet Dev 2005;15(2):153–62.

[93] Koh A, Murray H. Probing chromosome dynamics in Bacillus subtilis. Methods Mol Biol 2016;1431:91–108.

