## Appendix 1 - Software Installation Instructions.pdf for "Evaluation of chromosome structure modelling tools in bacteria"

### ShNeigh[1]

#### 1. Download link

Download link: https://github.com/fangzhen-li/ShNeigh

#### 2. Installation

Directly using the downloaded ShRec3D.m file, you need to install Matlab in advance.

#### 3. Run

Input file format (heatmapMatrix)

```
# Format of input Hi-C contact matrix
D_00    D_01    D_02    ...    D_0n
D_10    D_11    D_12    ...    D_1n
D_20    D_21    D_22    ...    D_2n
...     ...     ...     ...    ...
D_n0    D_n1    D_n2    ...    D_nn
```

Usage:

*[posMatrix,retVals]*=ShNeigh(heatmapMatrix,method_type)

method_type:

1 for ShNeigh1; 2 for ShNeigh2

Output:

posMatrix: N*4 matrix, the first column is a vector that marks valid genome loci, columns 2-4 are coordinates

retVal: a structure that contains some additional information

#### 4. Custom executable program

Although the code provided by this software can run in Matlab, the installation file for Matlab is too large. Here, standalone files that can be executed in Linux system are provided in Docker.

The modified ShNeigh1.m file code is as follows:

```
function ShNeigh1(inFile, outFile)
% Load the input file containing the normalized HiC matrix.
heatmapMatrix=load(inFile,'-ascii');
```

```
alpha_in = 1.0;
[rMat,selbin]=matFilter(heatmapMatrix);
selbinIdx=find(selbin);
n=size(rMat,1);
sig_in=n*0.023;
cover=(nnz(rMat)-n)/n/(n-1);
rho_in=max((1-cover)*sqrt(n),min(3,0.2*sqrt(n)));

FreqMat=rMat;
%Gaussian neighboring matrix,fixed alpha,fixed rho,fixed sigma
[error_term,XYZ]=shNeigh_knownParams(FreqMat,selbin,alpha_in,rho_in,sig_in,1);

posMatrix=[];
for i=1:size(XYZ,2)
    posMatrix= [posMatrix;selbinIdx(i),XYZ(1,i),XYZ(2,i),XYZ(3,i)];
end

dlmwrite(outFile,posMatrix,'delimiter','\t','precision','%.18f')
end
```

The modified ShNeigh2.m file code is as follows:

```
function ShNeigh2(inFile, outFile)
% Load the input file containing the normalized HiC matrix.
heatmapMatrix=load(inFile,'-ascii');
alpha_in = 1.0;
[rMat,selbin]=matFilter(heatmapMatrix);
selbinIdx=find(selbin);
n=size(rMat,1);
sig_in=n*0.023;
cover=(nnz(rMat)-n)/n/(n-1);
rho_in=max((1-cover)*sqrt(n),min(3,0.2*sqrt(n)));

FreqMat=rMat;
%Gaussian neighboring matrix,variable alpha,fixed sigma
fMpar = @(x)shNeigh_knownParams(FreqMat,selbin,x(1),x(2),sig_in,1);
[alpharho,fval,exitflag,output] = fminsearch(fMpar,[alpha_in
rho_in],optimset('MaxFunEvals',1000));
[error_term,XYZ]=shNeigh_knownParams(FreqMat,selbin,alpharho(1),alpharho(2),sig_in,1)
;

posMatrix=[];
for i=1:size(XYZ,2)
    posMatrix= [posMatrix;selbinIdx(i),XYZ(1,i),XYZ(2,i),XYZ(3,i)];
end

dlmwrite(outFile,posMatrix,'delimiter','\t','precision','%.18f')
end
```

These six files (FitFreq.m, freq2dist.m, matFilter.m, MDSneigh.m, neighborMatrix.m, shNeigh_knownParams.m) remain unchanged. Then use matlab's mcc tool to compile it into a separate executable file:

```
mcc -m ShNeigh1.m FitFreq.m freq2dist.m matFilter.m MDSneigh.m neighborMatrix.m shNeigh_knownParams.m
mcc -m ShNeigh2.m FitFreq.m freq2dist.m matFilter.m MDSneigh.m neighborMatrix.m shNeigh_knownParams.m
```

The obtained executable file can run normally on a Linux system with MATLAB Runtime 2019b installed.

`ShNeigh1 input_file output_file`

`ShNeigh2 input_file output_file`

If MATLAB Runtime 2019b is not installed on the target system, please install it from the following address: https://ssd.mathworks.com/supportfiles/downloads/R2019b/Release/9/deplonxa64/MATLAB_Runtime_R2019b_Update_9_glnxa64.zip

##### 4. Example

An example is placed in /3CModeler/example/ShNeigh

### FLAMINGO[2]

##### 1. Download link

Download link: https://github.com/wangjr03/FLAMINGO

##### 2. Installation

This test was completed using a process pipeline, so only a portion of the R package needs to be installed.

Requirement:

- R 3.6.0
- parallel, mgcv (1.8-28), Matrix (1.2-17), prodlim (2023.03.31), nlme (3.1-139)

##### 3. Run

Input file format:

```
# Format of input Hi-C contact matrix
D_00    D_01    D_02    ...    D_0n
D_10    D_11    D_12    ...    D_1n
D_20    D_21    D_22    ...    D_2n
```

```
...      ...      ...      ...      ...
D_n0     D_n1     D_n2     ...      D_nn
```

Description of scripts:

The original implementation process of this algorithm consists of eight parts, and after modification, five of them can be used to achieve it.

1. *1_preprocess_HiC.R*: This script uses the alpha = -0.25 exponential transformation to convert the Hi-C interaction frequency data into paired distance data, and the interaction frequency matrix and distance matrix are stored as. Rdata files for later use. Command line usage: *Rscript 1_preprocess_HiC.R <path to the folder containing the Hi-C data> <output path (define the working directory)>*
2. *2_generate_fragment_data.R*: This script divides the entire chromosome into large DNA domains,e.g. 100 kb. The interaction frequency matrices and distance matrices are generated for each domain by further dividing domains into small DNA fragments, i.e. 5 kb. The intra-domain interaction frequency matrices and distance matrices are used for the reconstruction of intra-domain structures. Command line usage: *Rscript 2_generate_fragment_data.R <path to the working directory> <chromosome name, i.e. chr1>*
3. *6_genereate_backbone_data.R*: This script processes low-resolution Hi-C data, i.e. 100 kb-resolution, for the reconstruction of the domain-level structures. Command line usage: *Rscript 6_genereate_backbone_data.R <path to the 100kb-resultion Hi-C data> <the working directory> <chromosome index>*
4. *7_reconstruct_within_cellline.R*: This step reconstructs the intra-domain structures from intra-domain distance matrices. It takes a pairwise distance matrix as input and returns the 3D coordinate matrix. *Rscript 7_reconstruct_within_cellline.R <fraction of basis> <penalization term of distance between adjacent small DNA fragments> <distance threshold to be penalized between adjacent small DNA fragments> <input pairwise distance matrix of large DNA fragments> <input pairwise distance matrix of large DNA fragments> <output file name> <output directory>*
5. *8_ensemble_structure_V2.R*: This script assembles the intra-domain structures with the domain-level structures by rotating the intra-domain structures. Command line usage: *Rscript 8_ensemble_structure.R <100kb interaction frequency matrix> <constructed backbone structure> <path to the output directory of step 7> <5kb pairwise distance matrix> <output file name>*

The output file is _coordinates.txt with three columns.

##### 4. Example

An example is placed in /3CModeler/example/FLAMINGO

### EVR$^3$

##### 1. Download link

Download link: https://github.com/mbglab/EVR

#### 2. Installation

Requirement:

- python (2.7 or 3.x)
- numpy
- scipy

Optinal:

- matplotlib (For plotting)
- Cython (Accelerating by multi-core CPUs)
- PyOpenCL (Accelerating by multi-core CPUs or many-core Processors like GPUs/APUs. Need to install related drivers.)

Speed up the program:

**With Cython**

The program uses Cython for acceleration by default. Iterative optimization code is written in Cython syntax (see CORE.pyx file). When running the program for the first time, you need to compile CORE.pyx into a dynamic link library by this command:

```
python setup.py build_ext --inplace
```

Before compiling, you need to install Cython by `pip install cython` and something alike. The build will produce a COREC.so file on a Linux system or a COREC.pyd file on a Windows system.

**With OpenCL**

In addition to using Cython for acceleration, you can use OpenCL. With OpenCL, you can speed up computing on CPUs and other processors like APUs and GPUs.

Firstly, you have to install the PyOpenCL library. On Linux, just type `pip install pyopencl` to install it. On Windows, you can download installation file on the (https://www.lfd.uci.edu/~gohlke/pythonlibs/#pyopencl), and use pip to install it (`pip install file_name`). In addition, you also need to install the relevant processor driver and OpenCL libraries. On Windows system, installation is easy. On Linux, you need to have administrator privileges and install the relevant OpenCL SDK.

To use OpenCL for acceleration, just add the "--OpenCL" parameter (no value):

```
python evr -i [Input file] -o [Output file] --OpenCL [...]
```

When using OpenCL to optimize the structure, the program will automatically detect the available OpenCL platforms. And then require the user to select the running platform and equipment (even if only one platform is detected). If you want to avoid user selection, you can add export PYOPENCL_CTX=id before the command. E.g:

```
export PYOPENCL_CTX='0'; python evr -i [Input file] -o [Output file] --OpenCL [...]
```

This command will make the program use the first platform detected.

##### 3. Run

```
python evr.py -i [input file] -o [output file] [...]
```

parameters_normal.txt:

```
-i: Input file
-o: Output file
-h: help messages
-se: Smoothing Factor, default = 2
-alpha: Factor in transforming interaction matrix into distance matrix, default = 0.5
--scale: Chromosome size, default = 20.0
--min_dis: Minimum distance between two bins, default = 0.001
--max_dis: maximum distance between two bins, default = 0.2
--iter_num: The number of iterations needed to run. If equal to 'auto', the program
will terminate when f value is less than 0.00001. default = 'auto'
-t: Number of threads, default = 'auto'
--seed: Seed for generating random initial structure, default = 'auto'
--OpenCL: Using OpenCL solver
--ChrType: Chromosome type. "0": Circular chromosome; "1": Linear chromosome. default
= '0'
```

Output file format:

The output file is a '.pdb' file that follows the standard PDB format. You can use the "plot.py" file to draw structure, or use any other PDB viewer program to do it.

```
python plot.py
/3CModeler/example/EVR/GSM2870407_mat_BC70_TACT_wt_MM_30C_normalized.pdb -o
/3CModeler/example/EVR/EVR/GSM2870407_mat_BC70_TACT_wt_MM_30C_normalized.pdf
```

##### 4. Example

An example is placed in /3CModeler/example/EVR

### PASTIS[4]

#### 1. Download link

Download link: https://github.com/hiclib/pastis

#### 2. Installation

Dependencies:

```
pip2 install numpy scipy scikit-learn pandas
```

Install PASTIS:

This package uses distutils, which is the default way of installing python modules.

```
python /3CModeler/tmp/pastis/setup.py install
```

This will install a python package pastis, and four programs: pastis-mds, pastis-nmds, pastis-pm1 and pastis-pm2. Calling any of those five programs will display the help.

#### 3. Run

Configure the config.ini file:

```
output_name: structure.pdb, str. Enter the file name, and the name of the algorithm
used will be added before the given file.
counts: data/counts.npy, str. Interaction matrix, npy format.
organism_structure: files/budding_structure.
resolution: 10000, integer. Data resolution.
chromosomes: "1, 2, 3, 4, 5, 6", str. Use comma to separate.
binary_mds: "MDS_all" MDS path.
binary_pm: "PM_all" PM path.
alpha: -3., float. Convert the interaction matrix to the coefficient of the expected
distance.
beta: 1., float. Structure scaling factor.
seed: 0, integer. Random seed with random structure at the beginning.
```

Input file format:

```
# Format of input Hi-C contact matrix(npy file)
D_00    D_01    D_02    ...     D_0n
D_10    D_11    D_12    ...     D_1n
D_20    D_21    D_22    ...     D_2n
...     ...     ...     ...     ...
D_n0    D_n1    D_n2    ...     D_nn
```

##### 4. Python file

The runPastis.py code is given in /3CModeler/scripts/pastis/ to customize the input and output.

Instructions:

```
python runPastis.py -w Working_directory -i InputFile -o OutputFile_Name -r Resolution
-a Alpha_of_IF_Matrix_to_Distance_Matrix -s Seed_of_initialization_structure
```

Parameter list:

```
-w: Working directory; default = "./", type = str
-o: Output File Name; type = str
-i: Input File; type = str, txt matrix format
-r: Resolution; default = 1000, type = int
-a: Alpha of IF Matrix to Distance Matrix; default = -1.0, type = float
-s: Seed of initialization structure; default = 0, type = int
```

### MOGEN[5]

##### 1. Download link

Download link: https://github.com/BDM-Lab/MOGEN

##### 2. Installation

Java installation required, please download from this address:

https://www.oracle.com/technetwork/java/javase/downloads/jdk8-downloads-2133151.html

You need to log in during the downloading process:

Username:

Password: Oracle123

After downloading, unzip and add environment variables:

```
# jdk
export JAVE_HOME="/3CModeler/thirdPart/jdk1.8.0_211"
export JRE_HOME="${JAVE_HOME}/jre"
export CLASSPATH=".:${JAVA_HOME}/lib:${JRE_HOME}/lib:$CLASSPATH"
export JAVA_PATH="${JAVA_HOME}/bin:${JRE_HOME}/bin"
export PATH="$PATH:${JAVA_PATH}"
```

```

```

##### 3. Run

```
java -jar 3DGenorator.jar parameters_normal.txt
```

parameters_normal.txt:

```


NUM: The number of generated structures
NBR_OF_CHR: The number of chromosomes
INTER_IF_THRESHOLD: The threshold of dyeing internal interaction
ADJACENT_DIST: The maximum distance between adjacent bins
CONTACT_DIST: Interaction distance
POS_MIN_DIST: The closest distance between bins
NEG_MAX_DIST_INTRA: The maximum distance between adjacent bins inside the dyeing
POS_MAX_DIST_WEIGHT_FILE: The smallest weight file between bins
POS_MIN_DIST_WEIGHT_FILE: The maximum weight file between bins
NEG_MIN_DIST_WEIGHT_FILE: The maximum distance weight file between adjacent bins
NEG_MAX_DIST_WEIGHT_FILE: The maximum distance weight file between adjacent bins
OUTPUT_FOLDER: Output path
INPUT_FILE: Input file
VERBOSE: Whether to print information during operation
LEARNING_RATE: Learning rate
MAX_ITERATION: Maximum number of iterations
```

Input file format:

```
# contact.txt
>bin1   >bin2   >frequency
```

##### 4. Example

An example is placed in /3CModeler/example/MOGEN

##### 5. Python file

The runMOGEN.py code is given in /3CModeler/scripts/MOGEN/ to customize the input and output.

Instructions:

```
python runMOGEN.py -w Working_directory -i InputFileName -o OutputFolder --NegMin 1.0
--NegMax 1700 --PosMin 1.0 --PosMax 1.0 --NUM The_number_of_generating_structure --
INTER_IF_THRESHOLD INTER_IF_THRESHOLD --ADJACENT_DIST ADJACENT_DIST --CONTACT_DIST
```

```
CONTACT_DIST --POS_MIN_DIST POS_MIN_DIST --NEG_MAX_DIST_INTRA NEG_MAX_DIST_INTRA --
LEARNING_RATE LEARNING_RATE --MAX_ITERATION MAX_ITERATION
```

Parameter list:

```
-w: Working directory, type = str, default = "./"
-i: Input File Name, type =str
-o: Output Folder, type = str
--NegMin: type = float, default = 1.0
--NegMax: type = float, default = 1700
--PosMin: type = float, default = 1.0
--PosMax: type = float, default = 1.0
--NUM: The number of generating structure, type = int, default = 1
--INTER_IF_THRESHOLD: INTER_IF_THRESHOLD, type = float, default = 0.0001
--ADJACENT_DIST: ADJACENT_DIST, type = float, default = 1.5
--CONTACT_DIST: CONTACT_DIST, type = float, default = 6.0
--POS_MIN_DIST: POS_MIN_DIST, type = float, default = 0.2
--NEG_MAX_DIST_INTRA: NEG_MAX_DIST_INTRA, type = int, default = 30
--LEARNING_RATE: LEARNING_RATE, type = float, default = 0.01
--MAX_ITERATION: MAX_ITERATION, type = int, default = 2000
```

### ShRec3D[6]

#### 1. Download link

Download link: https://sites.google.com/site/julienmozziconacci/home/softwares

#### 2. Installation

Directly using the downloaded ShRec3D.m file, you need to install Matlab in advance.

#### 3. Run

Input file format(dat file):

```
D_00    D_01    D_02    ...    D_0n
D_10    D_11    D_12    ...    D_1n
D_20    D_21    D_22    ...    D_2n
...     ...     ...     ...    ...
D_n0    D_n1    D_n2    ...    D_nn
```

Process:

- Modify the file name to data.dat
- Open Matlab, and open the ShRec3D.m script in matlab
- Modify the working directory of matlab to the directory where data.dat is located
- Run script

#### 4. Custom input and output

The software provides Matlab source code, but the code can only use data.dat as input and 3Dcoor.xyz as output by default. In order to customize the input and output, modify the code as follows:

```
% The input file contains the normalized HiC matrix.
% The output file contains the 3D coordinates.

function shrec3d(inFile, outFile)
% Load the input file containing the normalized HiC matrix.
Data2=load(inFile);

% Invert the HiC matrix by taking the reciprocal of each element.
Data=1./(Data2);

% Obtain the size of the matrix.
n = size(Data, 1);

% Perform the ShRec3D algorithm to generate 3D coordinates.
for k=1:n
    i2k = repmat(Data(:,k), 1, n);
    k2j = repmat(Data(k,:), n, 1);
    Data = min(Data, i2k+k2j);
end

% Calculate the center of the 3D coordinates.
center=zeros(n,1);
for i=1:n
    for j=1:n
    center(i)=center(i)+Data(i,j)^2-1/n*(Data(j,j:n)*Data(j:n,j));
    end
end

% Set any negative values in the center to zero.
center(center<0)=0;

% Calculate the square root of the center divided by n.
center=sqrt(center/n);

% Calculate the distance matrix based on the center coordinates.
distmat=1/2*(repmat(center.*center,1,n)+repmat((center.*center)',n,1)- Data.*Data);

% Perform eigenvalue decomposition to obtain the principal components.
[V,D]=eigs(distmat,4);

% Extract the eigenvalues.
lambda=diag(D);
```

```
% Calculate the 3D coordinates using the eigenvalues and eigenvectors.
XYZ=[V(:,1)*sqrt(lambda(1)) V(:,2)*sqrt(lambda(2)) V(:,3)*sqrt(lambda(3))];

% Scale the coordinates.
scale=100/max(max(XYZ));
XYZ=XYZ*scale;

% figure,plot3(XYZ(:,1),XYZ(:,2),XYZ(:,3));
% Save the 3D coordinates to the output file in XYZ format.
save(outFile,'XYZ','-ascii');
end
```

Then use matlab's mcc tool to compile it into a separate executable file: `mcc -m ShRec3D.m`

Please note that the compiled executable file can run normally on Linux systems with MATLAB Runtime 2019b installed.

`ShRec3D input_file output_file`

If MATLAB Runtime 2019b is not installed on the target system, please install it from the following address: https://ssd.mathworks.com/supportfiles/downloads/R2019b/Release/9/deplonxa64/MATLAB_Runtime_R2019b_Update_9_glnxa64.zip

##### 5. Example

An example is placed in /3CModeler/example/ShRec3D

### 3DMax[7]

##### 1. Download link

Download link: https://github.com/BDM-Lab/3DMax

##### 2. Installation

The installation is located in the /3CModeler/bin directory.

##### 3. Run

```
java -jar 3DMax.jar paramater.txt
```

parameter.txt content:

```
# number of structures will be generated
NUM = 1

OUTPUT_FOLDER = output/gm06990_hic

INPUT_FILE = input/gm06990_hic/IFList_Chr_19_1mb.txt

# factor used to convert IF into distance = 1 / (IF^factor)
CONVERT_FACTOR = 0.6

# set VERBOSE = true for information during optimization printed out
VERBOSE = true

# learning rate for the optimization process
LEARNING_RATE = 1

# if the program fails to generate structures, try to reduce this learning rate
MAX_ITERATION = 10000
```

Input file format:

```
# contact.txt
>bin1   >bin2   >frequency

# Square Matrix Input format
0.0000,0.0000,0.0281,0.0268,0.0227,0.0189,0.0143,...,0.0126
0.0000,0.0000,0.0000,0.0305,0.0272,0.0235,0.0184,...,0.0166
0.0281,0.0000,0.0000,0.0000,0.0326,0.0259,0.0197,...,0.0174
0.0268,0.0305,0.0000,0.0000,0.0000,0.0328,0.0234,...,0.0199
0.0227,0.0272,0.0326,0.0000,0.0000,0.0000,0.0185,...,0.0204
0.0189,0.0235,0.0259,0.0328,0.0000,0.0000,0.0000,...,0.0180
0.0143,0.0184,0.0197,0.0234,0.0185,0.0000,0.0000,...,0.0000
...    ...    ...    ...    ...    ...    ...    ...    ...
0.0126,0.0166 0.0174,0.0199,0.0204,0.0180,0.0000,...,0.0000
```

#### 4. Example

An example is placed in /3CModeler/example/3DMax

#### 5. Python file

The run3DMax.py code is given in /3CModeler/scripts/3DMax to customize the input and output.

Instructions:

```
python run3DMax.py -w Working directory -i Input File Name -o Output Folder --NUM The
_number_of_generating_structure --FA The_convert_factor --LEARNING_RATE Learning_rate
--MAX_ITERATION Max_iteration
```

Parameter list:

```
-w: Working directory, default = "./", type = str
-i: Input File Name, type =str
-o: Output Folder, type = str
--NUM: The number of generating structure, type = int, default = 1
--FA: The convert factor of transforming IF Matrix to distance matrix, type = float,
default = 0.5
--LEARNING_RATE: Learning_rate, type = float, default = 0.001
--MAX_ITERATION: Max_iteration, type = int, default = 10000
```

### MiniMDS[8]

#### 1. Download link

Download link: https://github.com/seqcode/miniMDS

#### 2. Installation

Install Python dependencies using:

```
pip install -r requirements.txt
```

The following optional dependencies can be installed manually (python3):

mayavi (http://docs.enthought.com/mayavi/mayavi/installation.html#installing-with-pip) (for plotting)

ImageMagick (https://www.imagemagick.org/script/index.php) (for creating gifs)

#### 3. Run

Intra-chromosomal miniMDS:

Run command：

```
python2 minimds.py [Hi-C BED path]
python2 minimds.py -o output file [Hi-C BED path]
```

Input file format:

```
# miniMDS uses intra- or inter-chromosomal BED files as input. Data must be normalized
prior to use (for example, using HiC-Pro).
# Format:
chrA    bin1_start  bin1_end    chrB    bin2_start  bin2_end
normalized_contact_frequency

# Example: chr22 intra-chromosomal data at 10-Kbp resolution:
chr22 16050000 16060000 chr22 16050000 16060000 12441.5189291
...
```

Inter-chromosomal miniMDS:

Run command:

```
python2 minimds_inter.py [prefix] [inter-chromosomal resolution] [intra-chromosomal
resolution]
```

Optional parameters:

```
# By default, minimds_inter.py uses all human chromosomes other than Y. You can
specify any number of chromosomes (in order) using the option -c.
# To perform inter-chromosomal analysis on chromosomes 1 and 2:
python minimds_inter.py -c 1 -c 2 data/GM12878_combined 1000000 10000
# You can specify a different number of autosomes using -C. To perform inter-
chromosomal analysis on all yeast autosomes:
python minimds_inter.py -C 16 my_yeast_dir 100000 10000
```

Input file format:

```
# The usage of minimds_inter.py is similar to minimds.py, however inter-chromosomal
files are required in addition to intra-chromosomal. To avoid entering filenames
separately for each chromosome, you must name
your files using a standard format.
#Intra-chromosomal format:
{prefix}_{ChrA}_{resolution}{kb or Mb}.bed

# Example:
GM12878_combined_22_100kb.bed

# Inter-chromosomal format:
{prefix}_{ChrA}_{ChrB}_{resolution}{kb or mb}.bed

# where A is before B in:
1 2 3 4 5 6 7 8 9 10 11 12 13 14 15 16 17 18 19 20 21 22 X
```

```
# Example:
GM12878_combined_21_22_100kb.bed
```

Plotting:

```
# Read a structure:
import data_tools
structure = data_tools.structure_from_file("GM12878_combined_22_100kb_structure.tsv")

# Create an interactive 3D plot in Mayavi. (Mayavi allows you to rotate the image and
save a view.)
import plotting
plotting.plot_structure_interactive(structure, color=(0,0.5,0.7), radius=0.01,
enrichments=my_enrichments)
```

##### 4. Example

An example is placed in /3CModeler/example/miniMDS

### SIMBA3D[9]

##### 1. Download link

Download link: https://github.com/nerettilab/SIMBA3D

##### 2. Installation

Optional Cython Build:

```
# To build cython replace the `setup.py` with `setup_cython.py` with the following
command
cp setup_cython.py setup.py

# Uninstall any previous simba3d installations
pip2 uninstall simba3d

# You will need to have cython and numpy installed before you try to build the code.
pip2 install cython
pip2 install numpy

# You may potentially need some additional c++ build tools. If you are using a managed
python installation like anaconda, then it should contain the build tools required for
the install. use `pip2 install .` to install simba3d with the cython.
# If it does not work, then you can revert to the pure python install.
```

```
cp setup_pure_python.py setup.py
pip2 uninstall simba3d
pip2 install .
```

Installation dependencies:

```
## Required Packages
* numpy - needed for scientic programing
* scipy - needed for the optimization tools and other scientic programing

## Recomended Packages
* matplotlib - needed if you want to plot the results in python
* simplejson - recommended over standard json packaged with python
```

Installation:

```
# After installing the python dependencies, one can install simba3d using distutils
with the following command
python2 setup.py --install

# If pip is installed, then it can be installed from this directory using
pip2 install .

# If you do not have root access you can do an install using the user scheme
pip2 install . --user

# Establish soft connection
ln -s /3CModeler/thirdPart/miniconda2/bin/simba3d /usr/bin/simba3d
```

#### 3. Run

After installing simba3d, the simba3d command should be available.

```
simba3d -i <json_tasklist_file> -t <number_of_threads>
```

Output file format:

```
# Currently, can create reports in the following output formats:
    '.npz'  zipped NumPy array file
    '.mat'  MatLab Mat-File file
    '.txt'  text file
    '.json' text file
# The default format is '.npz' and it is recommended to use this output because it is
easier to load into python and convert to some other format if desired. Of course, You
```

```
can use the following command to create pdb files for the result.
simba3d-convertion --ext_out .pdb ./*.json
```

##### 4. Example

An example is placed in /3CModeler/example/SIMBA3D

### LorDG[10]

##### 1. Download link

Download link: https://github.com/BDM-Lab/LorDG

##### 2. Installation

Java installation required, please download from this address:

https://www.oracle.com/technetwork/java/javase/downloads/jdk8-downloads-2133151.html

You need to log in during the downloading process:

Username:

Password: Oracle123

After downloading, unzip and add environment variables:

```
# jdk
export JAVE_HOME="/3CModeler/thirdPart/jdk1.8.0_211"
export JRE_HOME="${JAVE_HOME}/jre"
export CLASSPATH=".:${JAVA_HOME}/lib:${JRE_HOME}/lib:$CLASSPATH"
export JAVA_PATH="${JAVA_HOME}/bin:${JRE_HOME}/bin"
export PATH="$PATH:${JAVA_PATH}"
```

##### 3. Run

To run the tool, type:

```
java -jar 3DDistanceBaseLorentz.jar parameters.txt
```

Parameters are configured in the 'parameters.txt' file:

```
# number of structures will be generated
NUM = 1

OUTPUT_FOLDER = output/

# hi-C contact file, each line contains 3 numbers (separated by a space) of a contact,
position_1 position_2 interaction_frequencies
INPUT_FILE = input/yeast/poisson_chainDres25_List_noise10_1.txt

# the factor used to convert IF to distance, distance = 1/(IF^factor), when not
specified, the program will search for it in range [0.1, 3.0], step = 0.1
CONVERT_FACTOR = 1.1

# CHROMOSOME_LENGTH: remove it if there is only one chromosome. If there are multiple
chromosomes in the input data, specify number of points (or beads) of chromosomes in
the input data, separated by a comma. These numbers must be consistent with the input
data.
# CHROMOSOME_LENGTH =
229,241,197,190,179,169,157,145,124,135,133,132,98,89,83,81,79,77,57,62,36,36,153,29

# set VERBOSE = true for information during optimization printed out
VERBOSE = true

# learning rate for the optimization process, if optimization fails, try to reduce
this value
LEARNING_RATE = 1.0

# maximum number of iterations, the optimization may converge before this number
MAX_ITERATION = 10000
```

#### 4. Example

An example is placed in /3CModeler/example/LorDG

#### 5. Python file

The runLorDG.py code is given in /3CModeler/scripts/LorDG to customize the input and output.

Instructions:

```
python runLorDG.py -i Input_File_Name -o Output_Folder --NUM
The_number_of_generating_structure --LEARNING_RATE LEARNING_RATE -- MAX_ITERATION
MAX_ITERATION
```

Parameter list:

```
-i: Input File Name, type =str
```

```
-o: Output File Name, type = str
--NUM: The number of generating structure, type = int, default = 1
--LEARNING_RATE: LEARNING_RATE, type = float, default = 1
--MAX_ITERATION: MAX_ITERATION, type = int, default = 10000
```

Input file format:

```
# contact.txt
>bin1   >bin2   >frequency
```

### TADbit[11]

#### 1. Download link

Download link: https://github.com/3DGenomes/tadbit

#### 2. Installation

Execute directly in the downloaded directory:

```
python2 setup.py install
```

#### 3. Run

Requirement for importing data format:

```
        chrT_001    chrT_002    chrT_003    chrT_004    chrT_005    chrT_006
chrT_001    629 164 88  105 10  35
chrT_002    86  612 175 110 40  29
chrT_003    159 216 437 105 43  73
chrT_004    100 111 146 278 70  42
chrT_005    16  36  26  79  243 39
chrT_006    19  50  53  42  37  224
```

```
# Import the tadbit library
from pytadbit import Chromosome

# Initialize the chromosome structure and import the data
chrom = Chromosome(name = "E.coli")
chrom.add_experiment("GSM2870407_mat_BC70_TACT_wt_MM_30C", resolution = 5000, hic_data
```

```
= "GSM2870407_mat_BC70_TACT_wt_MM_30C.tsv")

# Find TAD
chrom.find_tad("GSM2870407_mat_BC70_TACT_wt_MM_30C")
chrom.experiments["GSM2870407_mat_BC70_TACT_wt_MM_30C"].tads

# Build a 3D model
CONFIG = {'dmel_01': {'kforce': 5,
                      'lowfreq': -0.7,
                      'lowrdist': 100,
                      'maxdist': 600,
                      'upfreq': 0.3
                      }
         }
models =
chrom.experiments["GSM2870407_mat_BC70_TACT_wt_MM_30C"].model_region(n_models=100,
n_keep=100, n_cpus=8, keep_all=True, config=CONFIG['dmel_01'])

# Save the model
models.define_best_models(100)
models[0].write_xyz_babel(filePath)
```

#### 4. Example

An example is placed in /3CModeler/example/TADbit

#### 5. Python file

The runTADbit.py code is given in /3CModeler/scripts/TADbit to customize the input and output.

Instructions:

```
python2 runTADbit.py -i [Input File Name] -o [Output File Name]
```

Parameter list:

```
-i: Input File Name, type =str
-o: Output File Name, type = str
--RESOLUTION: Resolution, type = int, default = 5000
--KFORCE: kforce, type = int, default = 5
--LOWFREQ: low frequency, type = int, default = -0.7
--LOWRDIST: min distance, type = int, default = 100
--MAXDIST: max distance, type = int, default = 600
--UPFREQ: up frequency, type = int, default = 0.3
--N_MODELS: Number of models, type = int, default = 100
--N_KEEP: Number of keep, type = int, default = 100
--N_CUPS: Number of cpus, type = int, default = 8
```

### GEM[12]

#### 1. Download link

Download link: https://github.com/mlcb-thu/GEM

#### 2. Installation

Directly put all the MATLAB script files in your MATLAB path, you need to install Matlab in advance.

#### 3. Run

Directly run the m-file **GEM.m** with parameters in the directory.

- Example

  do the following at the MATLAB command line:

  ```
  GEM('./HiC.txt', './loci.txt', 1E4, 4, 5E12, 0,-1)
  ```

- Parameters

  ```
  HiC_file: File name of Hi-C map.

  loci_file: File name of genomic loci.

  max_iter: Maximum number of iterations. Default is 1E4.

  M: Number of conformations. Default is 4.

  lambdaE: Coefficient of energy term. Default is 5E12.

  infer_latent: Whether to infer the latent function (1/0).

  input_sizepara: Packing density (bp/nm) of one fragment provided by user.
  Default is -1, which means using the estimated value by GEM.
  ```

- Input file

  File of Hi-C map: A N × N (N is the number of genomic loci) symmetric matrix separated by the table delimiter. The elements of it represent interaction frequencies of the Hi-C map. Example file: HiC.txt.

  File of genomic loci: A N × 1 matrix separated by the table delimiter. The elements of it represent the sequence position of the genomic loci. The fragement length between two genomic loci is equal to the resolution (e.g., 10kb, 50kb or 100kb) of the Hi-C data. Example file: loci.txt.

The example files (HiC.txt and loci.txt) contain normalized the Hi-C map and the genomic loci of chromosome 14 for 1Mb bins.

- Output information

  Final total cost, data cost, energy cost and inferred latent funtion (optional).

- Output file

  conformation[1-M].txt: The reconstructed chromatin conformation 1-M. Each conformation is a N × 3 matrix.

  proportions.txt: The corresponding weights of conformations (M × 1 matrix).

#### 4. Custom executable program

Although the code provided by this software can run in Matlab, the installation file for Matlab is too large. Here, standalone files that can be executed in Linux system are provided in Docker.

The modified GEM.m file code is as follows:

```
function []=GEM(HiC_file,loci_file,max_iter,M,lambdaE,infer_latent,input_sizepara)

% ----------------------- Overview ------------------------ %
% Main function for GEM                                      %
% --------------------------------------------------------- %
% HIC_file: File name of Hi-C map                            %
% loci_file: File name of genomic loci                       %
% max_iter: Maximum number of iterations                     %
% M: Number of conformations                                 %
% lambdaE: Energy coefficient                                %
% infer_latent: Whether to infer the latent function (1/0)  %
% input_sizepara: Packing density provided by user.          %
%              -1 means using the estimated value by GEM.  %
% --------------------------------------------------------- %

args = input('Enter HiC_file, loci_file, max_iter, M, lambdaE, infer_latent,
input_sizepara:', 's');
arguments = strsplit(args, ' ');
HiC_file = arguments{1};
loci_file = arguments{2};
max_iter = str2double(arguments{3});
M = str2double(arguments{4});
lambdaE = str2double(arguments{5});
infer_latent = str2double(arguments{6});
input_sizepara = str2double(arguments{7});

% ---------------------- Preproccess ---------------------- %
% Data Preprocess before modeling.                           %
% --------------------------------------------------------- %
% X: Hi-C map                                                %
% loci: Genomic loci                                         %
```

```
% sizepara: Packing density                                  %
% index: The index of the genomic loci of interest           %
% ---------------------------------------------------------- %
[ X,loci,sizepara,index ] = Preprocess(HiC_file,loci_file,input_sizepara);


% ---------------------- Optimizer ----------------------- %
% An optimization process that considers both Hi-C data     %
% and conformation energy.                                   %
% ---------------------------------------------------------- %
% structure: The reconstructed ensemble of conformations    %
%            (N*3*M matrix)                                  %
% proportions: The corresponding weights of conformations   %
% C: Total cost (C1+lambda_E*C2)                             %
% C1: Data cost (KL divergence)                              %
% C2: Energy cost (Conformation energy)                      %
% ---------------------------------------------------------- %
[structure,proportions,C,C1,C2]=Optimizer(X,loci,max_iter,sizepara,M,lambdaE,index);


disp('============================ RESULTS ============================');

%The conformations all have been aligned using the singular value decomposition (SVD)
algorithm
if M>1
    for i=1:M
        [ RMSE,structure(:,:,i) ] = SVD3D( structure(:,:,i),structure(:,:,1) );
    end
end
% Output reconstructed structure to 'conformation[1-M].txt'
for m=1:M
    dlmwrite(['conformation' num2str(m) '.txt'],structure(:,:,m),'delimiter',
'\t','precision','%6.4f');
end

% Output the corresponding weights of conformations to 'proportions.txt'
dlmwrite('proportions.txt',proportions,'delimiter', '\t','precision','%6.4f');

disp (['Total cost (C): ' num2str(C)]);
disp (['Data cost (C1): ' num2str(C1)]);
disp (['Energy cost (C2): ' num2str(C2)]);
disp (['The reconstructed conformations and the conformation weights are written to
''conformation[1-' num2str(M) '].txt'' and ''proportions.txt'', respectively. ' ]);

% ------------------- Latent function  -------------------- %
% Capture the latent function by comparing the modeled      %
% structures with the original Hi-C data.                    %
% ---------------------------------------------------------- %
if infer_latent==1
    LatentFunction( X(index,index),structure,M,proportions );
end

disp('=================================================================');
```

```
end
```

The remaining .m files remain unchanged. Then use matlab's mcc tool to compile it into a separate executable file:

```
mcc -m GEM.m BasicOptCell.m BayesianScore.m BayesParaSelect.m Energy.m GEMpara.m
LatentFunction.m Optimizer.m penalty.m Preprocess.m ResolutionMap.m RigidTransform3D.m
SVD3D.m Volume.m
```

The obtained executable file can run normally on a Linux system with MATLAB Runtime 2019b installed.

```
[root@5f1c3fa47c02 GEM]# GEM
Enter HiC_file, loci_file, max_iter, M, lambdaE, infer_latent,
input_sizepara:data/HiC.txt data/loci.txt 1E4 4 5E12 0 -1
......
```

##### 4. Example

An example is placed in /3CModeler/example/GEM

### NOTES

While AutoChrom3D, BACH, HSA, and MBO are not encompassed within this specific assessment, it is noteworthy that their corresponding packages have been made available in the /3CModeler/tmp/ directory, allowing readers the opportunity to peruse them.

### References


1. Li FZ, Liu ZE, Li XY, Bu LM, Bu HX, Liu H, Zhang CM. Chromatin 3D structure reconstruction with consideration of adjacency relationship among genomic loci. BMC Bioinform. 2020, 21(1):272, https://doi.org/10.1186/s12859-020-03612-4.
2. Wang H, Yang JX, Zhang Y, Qian JL, Wang JR. Reconstruct high-resolution 3D genome structures for diverse cell-types using FLAMINGO. Nat. Commun. 2022, 13(1):2645, https://doi.org/10.1038/s41467-022-30270-2.
3. Hua KJ, Ma BG. EVR: reconstruction of bacterial chromosome 3D structure models using error-vector resultant algorithm. BMC Genom. 2019, 20(1):738, https://doi.org/10.1186/s12864-019-6096-0.

4. Varoquaux N, Ay F, Noble WS, Vert JP. A statistical approach for inferring the 3D structure of the genome. Bioinformatics 2014, 30(12):i26-i33, https://doi.org/10.1093/bioinformatics/btu268.
5. Trieu T, Cheng J. MOGEN: a tool for reconstructing 3D models of genomes from chromosomal conformation capturing data. Bioinformatics 2016, 32(9):1286-1292, https://doi.org/10.1093/bioinformatics/btv754.
6. Lesne A, Riposo J, Roger P, Cournac A, Mozziconacci J. 3D genome reconstruction from chromosomal contacts. Nat. Methods. 2014, 11(11):1141-1143, https://doi.org/10.1038/nmeth.3104.
7. Oluwadare O, Zhang Y, Cheng J. A maximum likelihood algorithm for reconstructing 3D structures of human chromosomes from chromosomal contact data. BMC Genom. 2018, 19(1):161, https://doi.org/10.1186/s12864-018-4546-8.
8. Rieber L, Mahony S. miniMDS: 3D structural inference from high-resolution Hi-C data. Bioinformatics. 2017, 33(14):i261-i266, https://doi.org/10.1093/bioinformatics/btx271.
9. Rosenthal M, Bryner D, Huffer F, Evans S, Srivastava A, Neretti N. Bayesian Estimation of Three-Dimensional Chromosomal Structure from Single-Cell Hi-C Data. J. Comput. Biol. 2019, 26(11):1191-1202, https://doi.org/10.1089/cmb.2019.0100.
10. Trieu T, Cheng J. 3D genome structure modeling by Lorentzian objective function. Nucleic Acids Res. 2017, 45(3):1049-1058, https://doi.org/10.1093/nar/gkw1155.
11. Serra F, Baù D, Goodstadt M, Castillo D, Filion GJ, Marti-Renom MA. Automatic analysis and 3D-modelling of Hi-C data using TADbit reveals structural features of the fly chromatin colors. PLoS Comput. Biol. 2017, 13(7):e1005665, https://doi.org/10.1371/journal.pcbi.1005665.
12. Zhu G, Deng W, Hu H, Ma R, Zhang S, Yang J, Peng J, Kaplan T, Zeng J. Reconstructing spatial organizations of chromosomes through manifold learning. Nucleic Acids Res. 2018, 46(8):e50, https://doi.org/10.1371/10.1093/nar/gky065.
