## Appendix 2 - Untested Software.pdf for "Evaluation of chromosome structure modelling tools in bacteria"


### Supplementary information

### Untested software

| Software | Language | Sampling algorithm | Test result | Reason for not testing | Reference |
|---|---|---|---|---|---|
| Chromosome3D | Perl | Monte Carlo optimization using the Metropolis-Hastings algorithm with simulated annealing | No | Undisclosed installation dependencies | [1] |
| Chrom3D | Perl | Gibbs sampler with hybrid MC, and adaptive rejection sampling (ARS) | No | Missing input data information | [2] |
| HSA | R | GLM framework with Hamiltonian dynamics with simulated annealing | No | Download link is not accessible | [3] |
| PGS | Python | Simulated annealing / molecular dynamics | No | Unpublished code | [4] |
| tRex | R | Metropolis-Hastings algorithm / Gibbs sampler and Hamiltonian MCMC | No | The program cannot be downloaded | [5] |
| ChromSDE | Matlab | Linear and Quadratic Semi-definite programming (SDP) | No | The program cannot be downloaded | [6, 7] |
| ShRec3D+ | Matlab | Floyd-Warshall algorithm | No | Program not found | [8] |
| 5C3D | | Gradient descent | No | The program cannot be downloaded | [9] |
| AutoChrom3D | Perl | Non-linear constrained optimization | No | Not easy to install | [10] |
| GEM-FISH | Matlab | Gradient descent | No | FISH data missing | [11] |
| Gen3D | C++ | Adaptation, Simulated annealing and Genetic algorithm | No | Missing input file information | [12] |
| Hierarchical 3DGenome | Java | Gradient ascent and hierarchical modeling | No | Missing input file | [13] |
| ISDHiC | C, C++, Python | MCMC sampling using Hamiltonian MC | No | The program has an unknown error | [14] |
| MBO | Matlab | Manopt-manifold optimization | No | Single cell data required | [15] |
| MCMC5C | Java | Markov chain Monte Carlo (MCMC) sampling using the Metropolis-Hastings algorithm | No | The program cannot be downloaded | [16] |
| FisHiCal | R | SMACOF algorithm | No | Missing input data | [17] |
| InfMod3DGen | Matlab | Gradient ascent | No | The program runs incorrectly | [18] |

### References


[1] Adhikari, B.; Trieu, T.; Cheng, J. L. Chromosome3D: reconstructing three-dimensional chromosomal structures from Hi-C interaction frequency data using distance geometry simulated annealing. *BMC Genom.* **2016**, *17*, 886.

[2] Paulsen, J.; Sekelja, M.; Oldenburg, A. R.; Barateau, A.; Briand, N.; Delbarre, E.; Shah, A.; Sorensen, A. L.; Vigouroux, C.; Buendia, B.; Collas, P. Chrom3D: three-dimensional genome modeling from Hi-C and nuclear lamin-genome contacts. *Genome Biol.* **2017**, *18*, 21.

[3] Zou, C. C.; Zhang, Y. P.; Ouyang, Z. Q. HSA: integrating multi-track Hi-C data for genome-scale reconstruction of 3D chromatin structure. *Genome Biol.* **2016**, *17*, 40.

[4] Tjong, H.; Li, W. Y.; Kalhor, R.; Dai, C.; Hao, S. L.; Gong, K.; Zhou, Y. G.; Li, H. C.; Zhou, X. J.; Le Gros, M. A.; Larabell, C. A.; Chen, L.; Alber, F. Population-based 3D genome structure analysis reveals driving forces in spatial genome organization. *Proc. Natl. Acad. Sci. U.S.A.* **2016**, *113*, E1663-E1672.

[5] Park, J.; Lin, S. L. Impact of data resolution on three-dimensional structure inference methods. *BMC Bioinform.* **2016**, *17*, 70.

[6] Lin, D.; Hong, P.; Zhang, S. H.; Xu, W. Z.; Jamal, M.; Yan, K. J.; Lei, Y. Y.; Li, L.; Ruan, Y. J.; Fu, Z. F.; Li, G. L.; Cao, G. Digestion-ligation-only Hi-C is an efficient and cost-effective method for chromosome conformation capture. *Nat. Genet.* **2018**, *50*, 754-763.

[7] Zhang, Z.; Li, G.; Toh, K.-C.; Sung, W.-K. In *Inference of Spatial Organizations of Chromosomes Using Semi-definite Embedding Approach and Hi-C Data*, Research in Computational Molecular Biology, Berlin, Heidelberg, 2013; Deng, M.; Jiang, R.; Sun, F.; Zhang, X., Eds. Springer Berlin Heidelberg: Berlin, Heidelberg, 2013; pp 317-332.

[8] Li, J. G.; Zhang, W.; Li, X. D. 3D Genome Reconstruction with ShRec3D+and Hi-C Data. *IEEE/ACM Trans Comput Biol Bioinform* **2018**, *15*, 460-468.

[9] Fraser, J.; Rousseau, M.; Shenker, S.; Ferraiuolo, M. A.; Hayashizaki, Y.; Blanchette, M.; Dostie, J. Chromatin conformation signatures of cellular differentiation. *Genome Biol.* **2009**, *10*, R37.

[10] Peng, C.; Fu, L. Y.; Dong, P. F.; Deng, Z. L.; Li, J. X.; Wang, X. T.; Zhang, H. Y. The sequencing bias relaxed characteristics of Hi-C derived data and implications for chromatin 3D modeling. *Nucleic Acids Res.* **2013**, *41*, e183.

[11] Abbas, A.; He, X.; Niu, J.; Zhou, B.; Zhu, G. X.; Ma, T.; Song, J. P. K.; Gao, J. T.; Zhang, M. Q.; Zeng, J. Y. Integrating Hi-C and FISH data for modeling of the 3D organization of chromosomes. *Nat. Commun.* **2019**, *10*, 2049.

[12] Nowotny, J.; Ahmed, S.; Xu, L. F.; Oluwadare, O.; Chen, H.; Hensley, N.; Trieu, T.; Cao, R. Z.; Cheng, J. L. Iterative reconstruction of three-dimensional models of human chromosomes from chromosomal contact data. *BMC Bioinform.* **2015**, *16*, 338.

[13] Trieu, T.; Oluwadare, O.; Cheng, J. L. Hierarchical Reconstruction of High-Resolution 3D Models of Large Chromosomes. *Sci. Rep.* **2019**, *9*, 4971.

[14] Carstens, S.; Nilges, M.; Habeck, M. Inferential Structure Determination of Chromosomes from Single-Cell Hi-C Data. *PLoS Comp. Biol.* **2016**, *12*, e1005292.

[15] Paulsen, J.; Gramstad, O.; Collas, P. Manifold Based Optimization for Single-Cell 3D Genome Reconstruction. *PLoS Comp. Biol.* **2015**, *11*, e1004396.

[16] Rousseau, M.; Fraser, J.; Ferraiuolo, M. A.; Dostie, J.; Blanchette, M. Three-dimensional

modeling of chromatin structure from interaction frequency data using Markov chain Monte Carlo sampling. *BMC Bioinform.* **2011**, *12*, 414.

[17] Shavit, Y.; Hamey, F. K.; Lio, P. FisHiCal: an R package for iterative FISH-based calibration of Hi-C data. *Bioinformatics* **2014**, *30*, 3120-3122.

[18] Wang, S. Y.; Xu, J. B.; Zeng, J. Y. Inferential modeling of 3D chromatin structure. *Nucleic Acids Res.* **2015**, *43*, e54.
